# Differential neural mechanisms underlie cortical gating of visual spatial attention mediated by alpha-band oscillations

**DOI:** 10.1101/2023.08.21.553303

**Authors:** Xiaofang Yang, Ian C. Fiebelkorn, Ole Jensen, Robert T. Knight, Sabine Kastner

## Abstract

Selective attention relies on neural mechanisms that facilitate processing of behaviorally relevant sensory information while suppressing irrelevant information, consistently linked to alpha-band oscillations in human M/EEG studies. We analyzed cortical alpha responses from intracranial electrodes implanted in eight epilepsy patients, who performed a visual spatial attention task. Electrocorticographic data revealed a spatiotemporal dissociation between attention-modulated alpha desynchronization, associated with the enhancement of sensory processing, and alpha synchronization, associated with the suppression of sensory processing, during the cue-target interval. Dorsal intraparietal areas contralateral to the attended hemifield primarily exhibited a delayed and sustained alpha desynchronization, while ventrolateral extrastriatal areas ipsilateral to the attended hemifield primarily exhibited an earlier and sustained alpha synchronization. Analyses of cross-frequency coupling between alpha phase and broadband high-frequency activity (HFA) further revealed cross-frequency interactions along the visual hierarchy contralateral to the attended locations. Directionality analyses indicate that alpha phase in early and extrastriatal visual areas modulated HFA power in downstream visual areas, thus potentially facilitating the feedforward processing of an upcoming, spatially predictable target. In contrast, in areas ipsilateral to the attended locations, HFA power modulated local alpha phase in early and extrastriatal visual areas, with suppressed interareal interactions, potentially attenuating the processing of distractors. Our findings reveal divergent alpha-mediated neural mechanisms underlying target enhancement and distractor suppression during the deployment of spatial attention, reflecting enhanced functional connectivity at attended locations, while suppressed functional connectivity at unattended locations. The collective dynamics of these alpha-mediated neural mechanisms play complementary roles in the efficient gating of sensory information.

**SIGNIFICANCE STATEMENT:** Selective attention relies on neural mechanisms involved in target enhancement and distractor suppression to guide behavior. Using electrocorticographic data in humans, we show a spatiotemporal dissociation between cortical activities engaged in target facilitation and distractor inhibition during attentional deployment. We also found that, at attended locations, interareal interactions are enhanced through cross-frequency coupling along the visual hierarchy to potentially facilitate the processing of a spatially predictable target. In contrast, at unattended locations, intraareal interactions are enhanced through cross-frequency coupling, and interareal interactions are suppressed, together to potentially attenuate the processing of distractors. Our findings reveal that such a distributed cortical organization and complementary neural mechanisms enable efficient gating and filtering of sensory information in the anticipatory processing of spatial attention.

## INTRODUCTION

The selection of behaviorally relevant information from cluttered environments (i.e., selective attention) involves several neural mechanisms to facilitate the processing of behaviorally relevant sensory information, while suppressing that of irrelevant and potentially distracting information (1). In the non-human primate brain, modulation of rhythmic neural activity within the alpha frequency band (8-14 Hz) has been related to the selection process (2–4). Similarly, in the human brain, alpha-band activity has been closely associated with sustained attention to prioritize and filter incoming sensory information based on its behavioral relevance (5–9). Here, we investigated attentional modulation in the anticipatory processing of visual-spatial information, focusing on the complementary effects of alpha-band activity in selective attention and its role in orchestrating effective communication through cortical networks.

EEG and MEG studies suggest that decreases in alpha-band activity reflect the boosting of task-relevant sensory signals, while increases in alpha-band activity reflect active suppression of task-irrelevant sensory signals (8, 10, 11). For example, studies that manipulate visual spatial attention (e.g., with a spatially informative cue directing attention to a visual hemifield) have shown that alpha-band activity decreases at behaviorally relevant locations and increases at irrelevant or potentially distracting locations (5, 12–14). Such alpha lateralization has been found to correlate with behavioral performance during target detection, such that stronger alpha lateralization is predictive of faster responses and better target discriminability (15, 16). Studies have also shown a causal relationship between alpha-band activity and behavioral performance (i) by briefly disrupting posterior alpha oscillations using rhythmic transcranial magnetic stimulation (TMS), resulting in impaired target detection during spatial and feature-based attention (17–19), and (ii) by using MEG neurofeedback signals to manipulate parietal alpha lateralization, resulting in a spatial bias during online visual processing (20). Studies of non-human primates have further revealed a link between alpha oscillations and neuronal firing rates, suggesting that increased alpha-band activity is associated with a rhythmic inhibition of neuronal spiking (21, 22).

Most previous studies using EEG or MEG recordings to investigate sustained visual attention have found alpha desynchronization (associated with sensory enhancement) and synchronization (associated with sensory suppression) in posterior parietal and occipital regions (15, 23, 24). Attempts have been made to dissociate the temporal characteristics of posterior alpha desynchronization from synchronization in EEG, suggesting an early and transient alpha power decrease during shifting of spatial attention, followed by a later and sustained alpha power increase during maintenance of attention (25). Yet, the spatiotemporal dynamics of alpha-band activity at mesoscopic cortical level, and how it influences cortical excitability and functional interactions across the human visual processing hierarchy are not well understood. Electrocorticography (ECoG) provides superior spatial resolution from electrophysiological recordings obtained from subdural grid electrodes implanted in patients with intractable epilepsy. In addition, higher-frequency activity (∼70-200 Hz) is also obtained during ECoG recordings, and this broadband high-frequency activity (HFA) is strongly correlated with local cortical activation and spiking (26–31). ECoG recordings therefore provide a unique opportunity (i) to extend our understanding of the within-region and network-level dynamics associated with attention-modulated alpha-band activity in humans (32–35), and (ii) to advance our understanding of the relationship between population spiking activity and low-frequency neural oscillations (36–39) both within and between cortical regions.

Here, we analyzed cortical neural responses from intracranial electrodes covering occipital, parietal, temporal, and frontal cortex implanted in epilepsy patients, who performed an Eriksen Flanker task (i.e., a classic spatial attention task). We first characterized the spatial selectivity of cortical alpha-band activity, and then examined its cortical topography and temporal dynamics during attentional deployment. We report a spatiotemporal dissociation between attention-modulated alpha desynchronization and synchronization in preparation for an upcoming, spatially predictable target. Our findings reveal divergent alpha-mediated neural mechanisms underlying sensory gating during the anticipatory deployment of spatial attention, reflecting enhanced interareal interactions associated with target facilitation, while suppressed interareal interactions associated with distractor inhibition.

## RESULTS

We analyzed intracranial field potentials from 569 electrodes implanted over parietal, occipital, temporal, and frontal cortex in 8 pre-surgical epilepsy patients, who performed a classical spatial attention task. Electrodes localized in visual topographic areas were identified using our probabilistic atlas of the human visual system (40) (for electrode coverage of individual patients, see **Figure S1**). Electrodes in non-topographic cortex, analyzed for comparison purposes, were reconstructed using the Harvard-Oxford cortical parcellation based on anatomical markers (41) (for ROIs in each of the cortical areas, see **Figure S5A**).

### Spatial attention task and behavioral results

We probed a variant of an Eriksen flanker task (42) that we also use in comparative electrophysiology studies in non-human primates (43) and human MEG studies (44, 45). Each trial started with a central fixation point presented on a computer monitor, followed by a brief spatial cue presented in one of 8-16 pseudo-randomized peripheral locations. After a variable delay interval, a circular array of barrel and bowtie shapes was presented, and patients indicated with a button press, which shape appeared at the cued location (**Figure 1A**). Either congruent or incongruent shape stimuli could flank the target shape, which is known to influence behavioral outcome (i.e., accuracy and reaction times, RTs). Patients completed between 150 and 300 trials of the task and achieved high accuracies ranging from 83 to 98% (mean ± SD = 93 ± 5%), including better accuracies in trials with congruent flanker stimuli (hit rate mean ± SD: congruent = 95 ± 4%, incongruent = 89 ± 7%; congruent > incongruent, *p* = 0.03, t-test) and suggesting that the patients performed the task as instructed. Because response speed was not emphasized in our task, RTs were not reliable measures of flanker effects. Only trials with correct responses with RTs ≤ 3 SD from the mean (within subject) were included for the analyses of neural data (median = 91.7% of total trials, min = 80.5%, max = 95.7%). Our analyses were focused on neural data obtained during the cue-target interval, i.e., during attentional deployment in anticipation of the target array. Details of demographic information and behavioral performance for each patient are available in (46).

**Figure 1.**
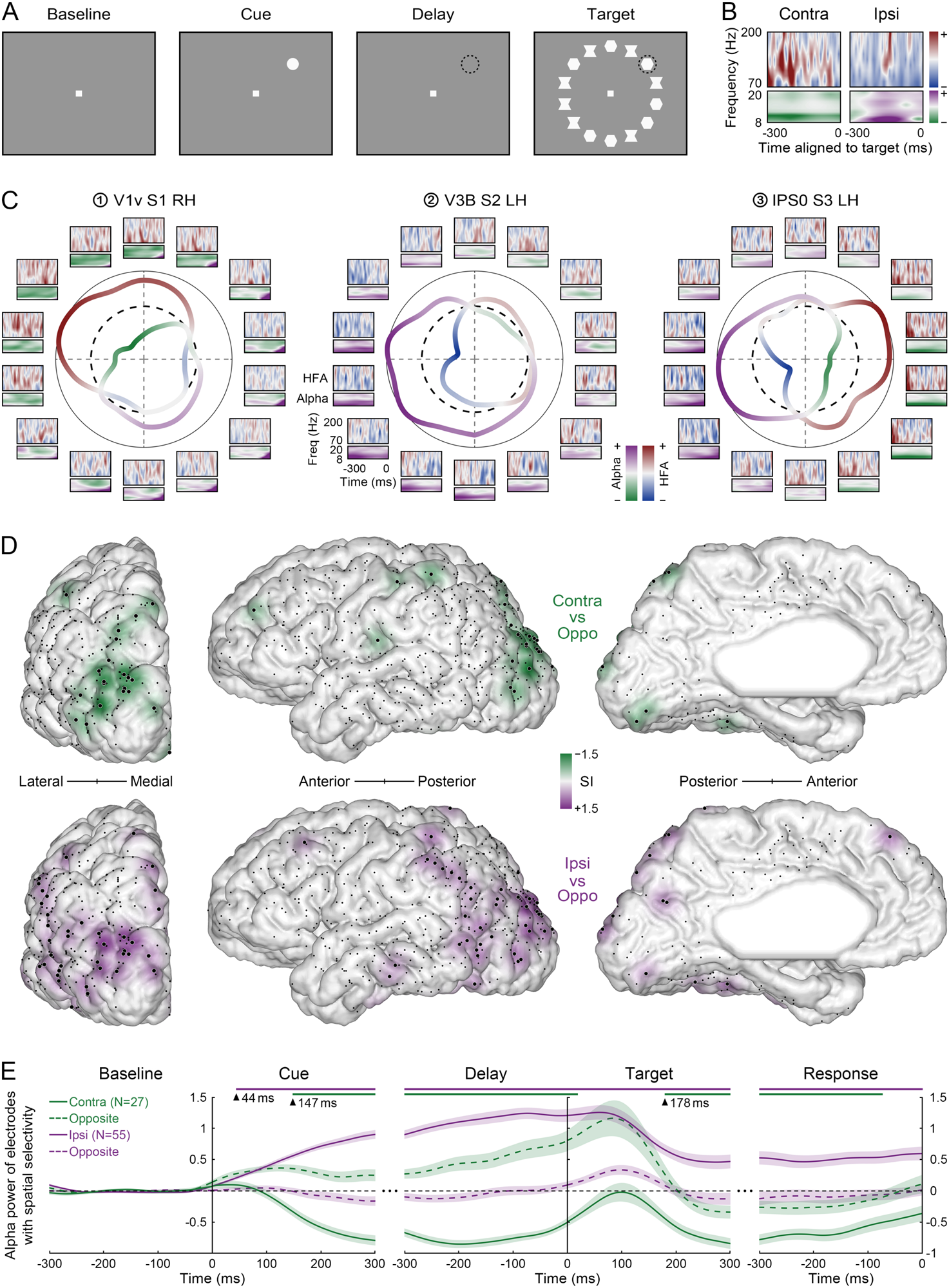
Experimental task and attentional modulation of alpha power. **A:** Each trial started with central fixation for 1100 ms followed by a brief spatial cue flashed for 100 ms. After a variable delay of 300-700 ms, a circular array of targets (barrel or bow tie shapes) was presented, and the subjects were instructed to detect the target shape at the cued location. **B:** Single-trial power spectra (normalized to baseline) are illustrated for the alpha-band activity (8-20 Hz, lower panels) and broadband high-frequency activity (HFA, 70-200 Hz, upper panels) at a posterior parietal electrode during the cue-target interval aligned to target onset, when attention was directed to the contralateral (left panels) or ipsilateral (right panels) visual hemifield. **C:** Response profiles are shown for three example electrodes in subjects S1-S3 respectively, recorded in the right hemisphere from early visual cortex V1v (1), and in the left hemisphere from extrastriate cortex V3B (2) and intraparietal sulcus area IPS0 (3). Power spectra (normalized to baseline and averaged across trials) are shown for the alpha-band activity (lower panels) and HFA responses (upper panels) during the cue-target interval aligned to target onset at each of the 14 cued locations. Central panels: smoothed spatial tuning curves illustrate normalized alpha power and HFA power averaged during the delay at each location. Black circles indicate normalized baseline (dashed line) and maximum response (solid line). **D:** Cortical topography of alpha RF_desynch_ and RF_synch_ SI values from electrodes that showed spatially selective alpha responses (large dots) during the cue-target interval, rendered across all subjects onto a standard brain surface shown from the posterior, lateral, and medial views (black dots denote cortical localization of all electrodes). Attention-modulated alpha desynchronization (alpha power at contralateral RF_desynch_ vs. opposite visual field locations) is illustrated in green showing a dorsal distribution (upper panels), whereas alpha synchronization (ipsilateral RF_synch_ vs. opposite locations) is illustrated in purple showing a ventrolateral distribution (lower panels). **E:** Time course of alpha power averaged across electrodes that showed spatially selective alpha responses (large dots in **D**) at contralateral RF_desynch_ (green solid line, N = 27) and ipsilateral RF_synch_ (purple solid line, N = 55) locations, in contrast with the corresponding opposite visual field locations (colored dashed lines) throughout the entire trial (shaded regions denote SEM). Lines above curves indicate time points of spatially selective alpha responses for the solid-line time series, when alpha power at RF_desynch_ or RF_synch_ were significantly different than the baseline (black dashed line) and the corresponding opposite locations (t-tests, p < 0.05, Bonferroni corrected); arrows denote the onset of significant time periods.

### Cortical alpha responses show local attention-modulated spatial selectivity

We have previously reported high spatial selectivity for broadband high-frequency activity (HFA, 70-200 Hz) in visual topographic cortex (46). Here, we measured the spatial selectivity of alpha-band activity (8-20 Hz) while spatial attention was allocated at one of 8-16 peripheral locations during the cue-target interval. One hundred and twenty-four electrodes were analyzed in topographically organized visual areas from five subjects, including early visual cortex (EVC: *V1-V3 dorsal/ventral*; *N* = 37), extrastriate cortex (ESC: *V3A/B, TO1-2, LO1-2 & VO1-2*; *N* = 42), and parietal cortex, particularly areas along the intraparietal sulcus (IPS: *IPS0-5 & SPL*; *N* = 45). Electrodes located outside of topographic cortex were broadly localized by lobe: parietal (*N* = 196), temporal (*N* = 131), and frontal (*N* = 118).

**Figure 1C** illustrates time-frequency profiles of three representative electrodes in visual topographic cortex, recorded from three subjects (S1-S3) and averaged across trials. These electrodes were located in visual areas *V1v* (right), *V3B* (left), and parietal area *IPS0* (left) respectively. We observed distinct response profiles during the cue-target interval with spatially tuned responses in both alpha-band activity and HFA responses. In agreement with previous M/EEG studies on visual attention (5, 8, 14), we observed a decrease of cortical alpha power relative to the pre-cue baseline when attention was directed to the contralateral visual hemifield, and an alpha power increase when attention was directed to the ipsilateral visual hemifield (single-trial responses illustrated in **Figure 1B**). This pattern of responses is consistent with the notion that alpha-band activity is associated with the suppression of visual processing (8, 10, 11).

To further investigate the cortical distribution of delay-related alpha responses, we first identified task-responsive electrodes that showed (i) a significant alpha power decrease (i.e., desynchronization) during the cue-target interval (300 ms before target onset), relative to the pre-cue baseline (300 ms before cue onset) (*p* < 0.05, permutation test), at contralateral attended locations, and (ii) a significant alpha power increase (i.e., synchronization) at ipsilateral unattended locations. These alpha power changes defined alpha desynchronization ECoG response fields (RF_desynch_) and alpha synchronization response fields (RF_synch_), respectively. For instance, in **Figure 1C-3**, we illustrate an example electrode in left IPS (*IPS0*), where the green curve indicates alpha RF_desynch_ in the right visual field and the purple curve indicates alpha RF_synch_ in the left visual field. Note that the alpha RF_desynch_ overlaps with locations that showed an increase of HFA power, whereas the alpha RF_synch_ overlaps with locations that showed a decrease of HFA power. At the population level, we found a small group of electrodes that showed both attention-related alpha desynchronization and synchronization effects (in total, *N* = 17, accounting for 3% of all electrodes; detailed breakdown in **Figure S2** & **Table S1**).

We next identified electrodes that only showed either attention-related alpha desynchronization, or synchronization. For instance, in **Figure 1C-1**, we illustrate an example electrode in right ventral EVC (*V1v*) where the green curve indicates alpha RF_desynch_ in the left upper visual field, overlapping with locations that showed an increase of HFA power. We further quantified the spatial selectivity of attention-modulated alpha desynchronization by measuring alpha power differences at RF_desynch_ locations as compared to opposite locations, referred to as alpha selectivity indices (SI). Electrodes with alpha RF_desynch_ SI values significantly less than the estimated SI distribution (< 95% CI) from non-responsive electrodes were considered showing attention-modulated spatial selectivity (*p* < 0.05). We found that visual topographic cortex had a higher percentage of electrodes than non-topographic cortex exhibiting attention-modulated spatially selective alpha desynchronization, particularly in IPS (*N* = 9, 20%), EVC (*N* = 5, 14%), and ESC (*N* = 5, 12%) (for comparison in non-topographic areas: parietal cortex - *N* = 4, 2%, frontal cortex - *N* = 2, 2%, and temporal cortex - *N* = 2, 2%) (**Figure S2** ‘RF_d_ Selective’ dark green dots, **Table S1** ‘*α_-_* Selective’ electrodes).

We also identified electrodes that only showed attention-related alpha synchronization. For instance, in **Figure 1C-2**, we illustrate an example electrode in left ESC (*V3B*) with an alpha RF_synch_ in the left visual field (purple curve), overlapping with locations that showed a decrease of HFA power. To further quantify the spatial selectivity of attention-modulated alpha synchronization, we measured alpha power differences at RF_synch_ locations as compared to opposite locations and identified electrodes with alpha RF_synch_ SI values significantly greater than the estimated SI distribution (> 95% CI) from non-responsive electrodes (*p* < 0.05). We found that visual topographic cortex also had a higher percentage of electrodes than non-topographic cortex exhibiting attention-modulated spatially selective alpha synchronization, particularly in ESC (*N* = 12, 29%), EVC (*N* = 6, 16%), and IPS (*N* = 3, 7%) (for comparison in non-topographic areas: parietal cortex - *N* = 23, 12%, temporal cortex – *N* = 9, 7%, and frontal cortex - *N* = 2, 2%) (**Figure S2** ‘RF_s_ Selective’ dark purple dots, **Table S1** ‘*α_+_* Selective’ electrodes). Taken together, these results suggest that recording sites with spatially selective alpha desynchronization contralateral to the cued location were more prominent in dorsal intraparietal areas, whereas recording sites with spatially selective alpha synchronization ipsilateral to the cued location were more prominent in ventrolateral extrastriatal areas. This pattern of responses is consistent with the notion that alpha-band activity is associated with retinotopic maps (45, 47), collectively revealing distinct cortical sources of attention-modulated spatially selective alpha desynchronization and synchronization.

**Figure 2.**
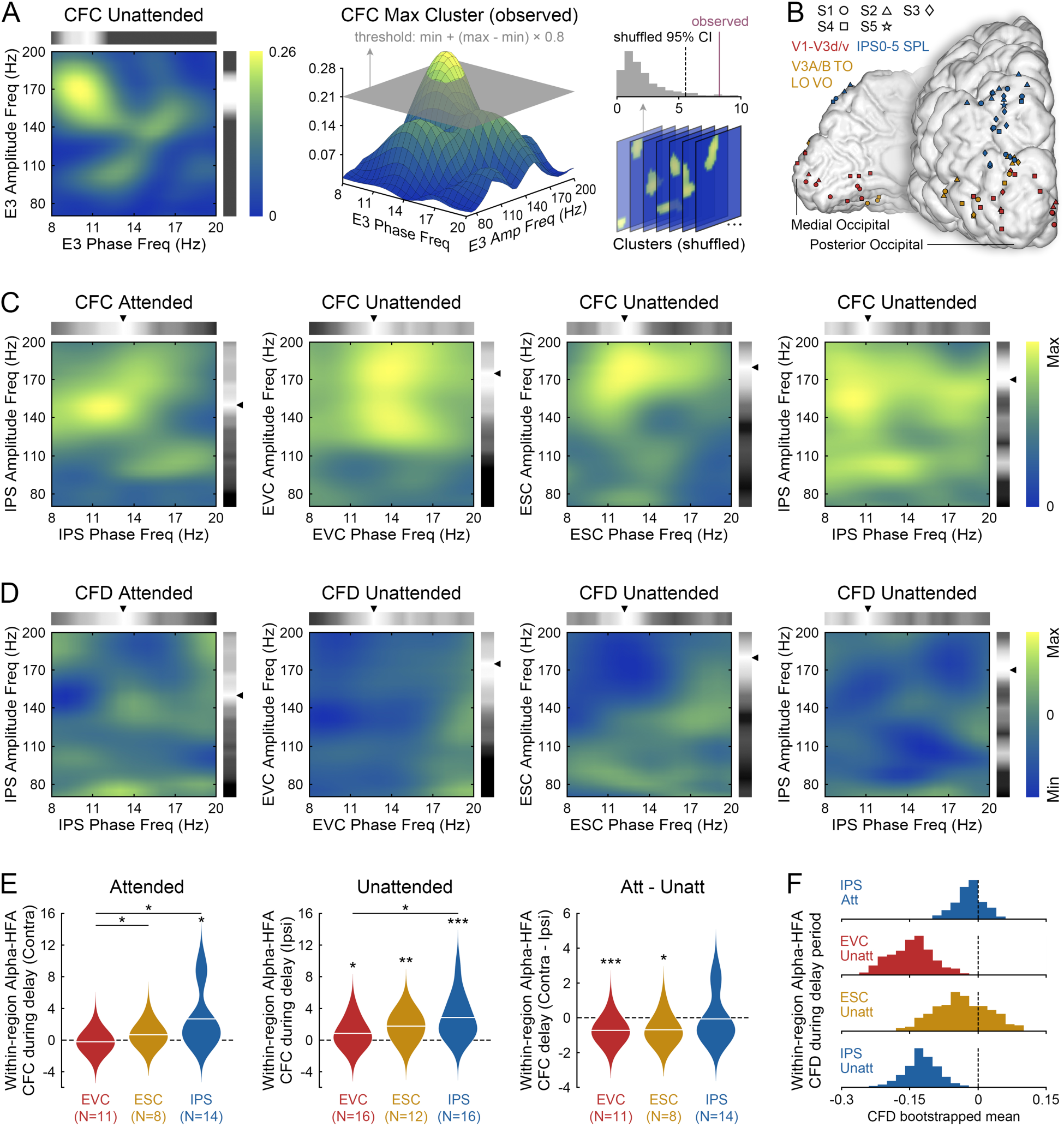
Attentional modulation of cross-frequency interactions within cortical regions. **A:** Within-region cross-frequency coupling (CFC) is illustrated for the example electrode E3 (**Figure 1C-3**) at unattended locations, highlighting the peak frequencies of alpha phase (upside horizontal bar) and HFA amplitude (right-side vertical bar) that contributed to the most dominant CFC effect (left panel). Electrodes with an observed CFC value (sum of the maximum CFC cluster exceeding threshold, middle panel) greater than the 95% CI of a shuffled surrogate distribution (right panel) were considered to demonstrate significant within-region CFC (p < 0.05). **B:** Electrodes that showed significant within-region CFC, during the delay at attended or unattended locations, in visual topographic cortex are color-coded for EVC (red), ESC (yellow), and IPS (blue). Electrodes are projected on a standard brain surface shown from the posterior and medial views, contributed from all five subjects (S1-S5 marked in different shapes) with coverage of visual topographic cortex. **C-D:** Within-region mean CFC (**C**) and cross-frequency directionality (CFD; **D**) are averaged across significant electrodes and illustrated for visual topographic areas that showed significant group-level increases of within-region CFC during the delay at attended and unattended locations (CFC phase-amplitude peak frequencies marked by arrows). **E:** Group-level within-region CFC during the cue-target delay at attended locations (left panel) and unattended locations (middle panel), normalized to the baseline-level CFC (dashed line) for each visual topographic area (numbers of contributing electrodes enclosed in parentheses). The attentional modulation effect of within-region CFC was estimated by contrasting group-level CFC values at attended locations with those at unattended locations (right panel) (t-tests statistical significance: * p < 0.05, ** p < 0.01, *** p < 0.001). **F:** Group-level mean directionality of within-region CFC was estimated through bootstrap iterations over CFD values from significant electrodes within each area at attended and unattended locations. The distributions of the bootstrapped CFD means are illustrated and evaluated against zero, with positive mean CFD indicating that alpha phase modulates HFA power, while negative mean CFD indicating that HFA power regulates alpha phase.

### Electrocorticographic data reveal spatiotemporal dissociation between attention-modulated alpha desynchronization and synchronization

To visualize the cortical topography of attention-modulated spatially selective alpha desynchronization and synchronization, we rendered all significant alpha RF_desynch_ and RF_synch_ SI values onto a standard brain surface (**Figure 1D**). The topography was spread cortically using a Gaussian kernel with colors indicating the magnitude of alpha spatial selectivity during attentional deployment. We observed a dorsal distribution of attention-modulated alpha desynchronization, covering electrodes over occipital visual cortex and dorsal intraparietal areas, when attention was directed to the contralateral visual field. In contrast, we observed a ventrolateral distribution of attention-modulated alpha synchronization, covering electrodes over occipital visual cortex and ventrolateral extrastriatal areas, when attention was directed to the ipsilateral visual field (alpha RF_desynch_ SI in IPS < 0: *p* = 7×10^−5^, RF_synch_ SI in ESC > 0: *p* = 1×10^−5^, t-tests; detailed statistics reported in **Figure S3**). We further examined the temporal dynamics of attention-modulated alpha cortical topography by computing alpha SI values within consecutive 100 ms time windows and observed a build-up of and sustained dorsal/ventrolateral distribution after the presentation of the spatial cue (300 ms after cue onset) and during the maintenance of spatial attention (300 ms before target onset) (**Figure S4**). Our findings suggest a cortical dissociation between alpha-band desynchronization and synchronization in preparation for an upcoming, spatially predictable target (i.e., during the cue-target interval), with (i) dorsal intraparietal areas primarily demonstrating attention-modulated desynchronization of alpha-band activity contralateral to the expected target location (i.e., the attended location) and (ii) ventrolateral extrastriatal areas primarily demonstrating attention-modulated synchronization of alpha-band activity ipsilateral to the expected target location.

To further analyze the temporal dynamics of attention-modulated alpha desynchronization and synchronization, we illustrated the time course of alpha power averaged across electrodes that showed spatially selective alpha responses throughout the entire trial (**Figure 1E**). We identified time periods when alpha power at RF_desynch_ or RF_synch_ were significantly different than the baseline and the corresponding opposite locations (*N* = 27 for the RF_desynch_ group and *N* = 55 for the RF_synch_ group, t-tests, *p* < 0.05, Bonferroni corrected for multiple comparisons). We observed a delayed onset of attention-modulated alpha desynchronization (147 ms after cue onset) that was then sustained during the cue-target interval and returned to the baseline level only, when attention was directed to the contralateral visual field. This desynchronization effect was then reestablished after the target array appeared (178 ms after target onset) and was maintained until the end of the trial. In contrast, we observed an early onset of attention-modulated alpha synchronization (44 ms after cue onset, approximately one alpha cycle earlier than the desynchronization onset) that was sustained throughout the entire trial when attention was directed to the ipsilateral visual field. Thus, our results also reveal a temporal dissociation between attention-modulated alpha desynchronization and synchronization, with (i) a delayed and sustained desynchronization of alpha-band activity during attention deployment contralateral to the expected target location (i.e., the attended location), followed by a reestablished and maintained desynchronization of alpha-band activity during target selection, and (ii) an earlier and sustained synchronization of alpha-band activity ipsilateral to the expected target location throughout the entire task trial.

### High-frequency population activity regulates local alpha phase at unattended locations

In sample electrodes (**Figure 1C**), we observed that maximal HFA responses co-occurred with maximal alpha desynchronization, whereas maximal HFA suppression co-occurred with maximal alpha synchronization, suggesting a potential relationship between HFA and alpha-band activity. Therefore, we probed whether HFA and alpha responses were independent of each other or, alternatively, influenced each other. Specifically, we examined whether the HFA power envelope (a proxy for population neuronal spiking (26, 27, 48) or superficial dendritic processes (31, 49)) was related to the phase of local alpha-band activity as previously found in monkey laminar activity (39) and in human MEG and ECoG activity (37, 50). We first measured electrode-specific cross-frequency coupling (CFC) between alpha phase and HFA amplitude during the cue-target interval, when the cue occurred at locations associated with maximal alpha desynchronization (contralateral attended locations - RF_desynch_) or maximal alpha synchronization (ipsilateral unattended locations - RF_synch_), focusing on electrodes that were localized in visual topographic cortex.

For instance, in the example electrode E3 (**Figure 1C-3**), we visualized the spectral features of within-region CFC at unattended locations, illustrating the peak frequencies of alpha phase and HFA amplitude that contributed to the most dominant phase-amplitude coupling at this electrode (**Figure 2A** left). In order to quantify the dominant coupling effect with consideration of spectral variance across electrodes, we set an electrode-specific threshold to identify the maximum cluster of CFC estimates exceeding the threshold, then calculated the sum of the maximum cluster referred to as CFC value for subsequent analyses (**Figure 2A** middle). Electrodes with an observed CFC value greater than the 95% confidence interval of a shuffled surrogate distribution were considered to demonstrate significant within-region CFC effect (*p* < 0.05) (**Figure 2A** right). Following this approach, we identified electrodes with significant within-region coupling between alpha phase and HFA amplitude at attended locations in visual topographic cortex, including EVC (*N* = 13, 35%), IPS (*N* = 14, 31%), and ESC (*N* = 9, 21%) (**Table S1** ‘*α_-_* Coupled’ electrodes). We also found electrodes that showed significant within-region CFC at unattended locations in visual topographic cortex, including EVC (*N* = 17, 46%), IPS (*N* = 16, 36%), and ESC (*N* = 14, 33%) (**Table S1** ‘*α_+_* Coupled’ electrodes). This within-region cross-frequency coupling effect was found in all five subjects (S1-S5), who had coverage of visual topographic cortex (**Figure 2B**).

**Figure 3.**
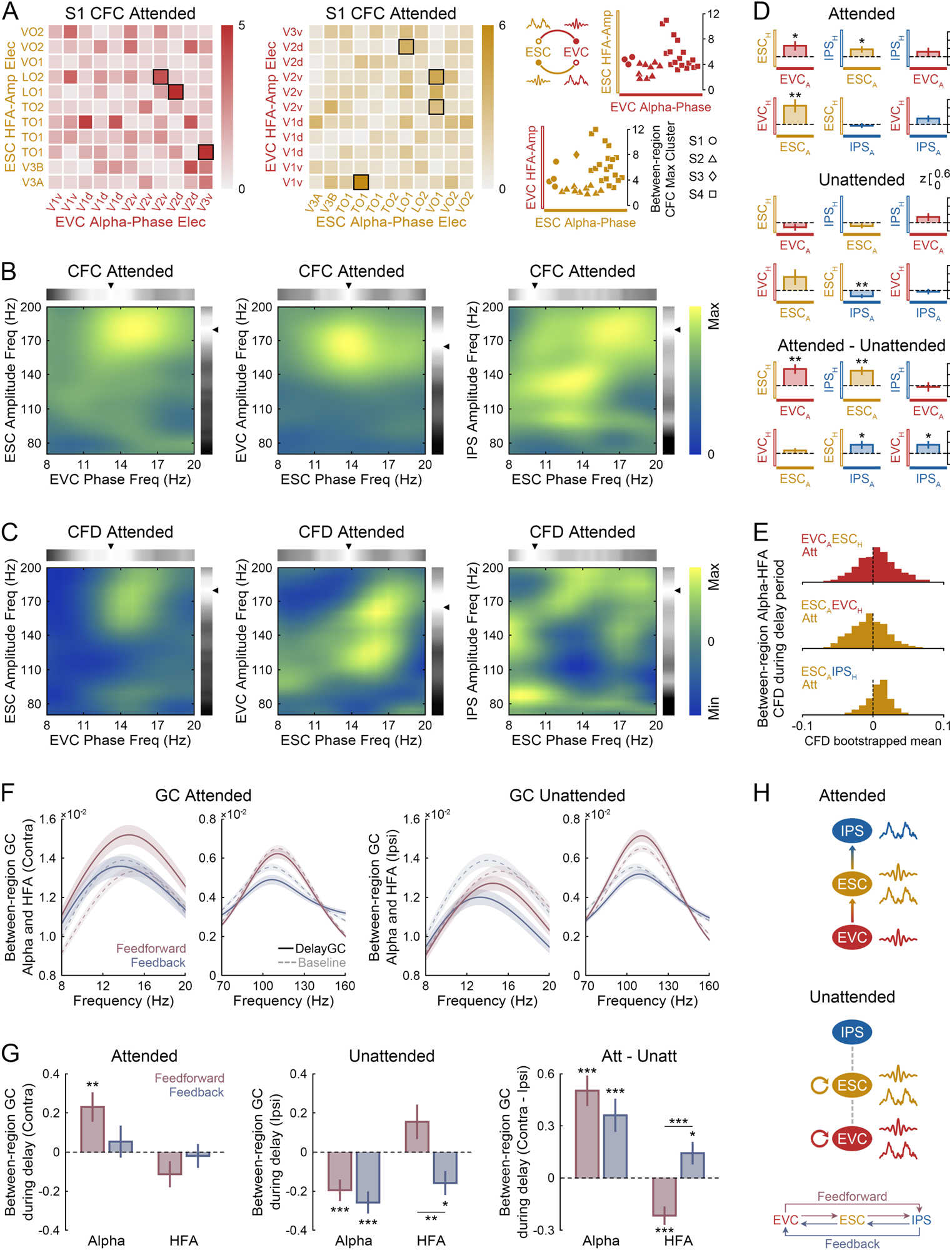
Attentional modulation of functional connectivity between cortical regions. **A:** Between-region cross-frequency coupling (CFC) is illustrated for the example pair of visual topographic areas between EVC (red) and ESC (yellow) at attended locations in subject S1, with alpha phase estimated from EVC electrodes and HFA amplitude from ESC electrodes (EVC_Alpha_-ESC_HFA_, left panel), as well as the reversed measure (ESC_Alpha_-EVC_HFA_, middle panel). Electrode pairs with a CFC maximum cluster greater than the 95% CI of a shuffled surrogate distribution were considered to demonstrate significant between-region CFC (p < 0.05) (outlined with black squares). These significant electrode pairs were identified in all four subjects (S1-S4 marked in different shapes) with multiregional coverage of visual topographic areas to estimate the group-level between-region CFC for each phase-amplitude measure within each pair of visual topographic areas (right panel). **B-C:** Between-region mean CFC (**B**) and CFD (**C**) are averaged across significant electrode pairs and illustrated for pairs of visual topographic areas that showed significant group-level increases of between-region CFC during the delay at attended locations (CFC phase-amplitude peak frequencies marked by arrows). **D:** Group-level between-region CFC during the cue-target interval at attended locations (upper panel) and unattended locations (middle panel), normalized to the baseline-level CFC (dashed line) for each pair of visual topographic areas. The attentional modulation effect of between-region CFC was estimated by contrasting group-level CFC values at attended locations with those at unattended locations (lower panel). **E:** Group-level mean directionality of between-region CFC was estimated through bootstrap iterations over CFD values from significant electrode pairs for each pair of visual topographic areas at attended locations. The distributions of the bootstrapped CFD means are illustrated and evaluated against zero, with positive mean CFD indicating that alpha phase modulates HFA power, while negative mean CFD indicating that HFA power regulates alpha phase. **F:** Between-region Granger causal (GC) influences of information flow along the visual hierarchy are illustrated for the feedforward and feedback pathways at attended locations (left panel) and unattended locations (right panel) separately for the alpha-band and HFA frequencies, averaged across significant electrode pairs during the cue-target interval (solid line) and the pre-cue baseline (dashed line) (shaded regions denote SEM). **G:** Group-level between-region GC influences during the cue-target interval at attended locations (left panel) and unattended locations (middle panel), normalized to the baseline-level GC (dashed line) for the feedforward and feedback GC influences in the alpha-band and HFA frequencies. The attentional modulation effect of between-region GC influences was estimated by contrasting group-level GC values at attended locations with those at unattended locations (right panel). All statistical significance was established using t-tests (* p < 0.05, ** p < 0.01, *** p < 0.001). **H:** Schematic diagram of intraareal and interareal cross-frequency interactions at attended locations (upper panel) and unattended locations (lower panel).

We next examined the effect of attentional deployment on the group-level within-region cross-frequency coupling by comparing CFC values from significant electrodes during the cue-target interval to the baseline-level CFC (combining baselines at attended and unattended locations) within each area. At attended locations, we found a significant group-level increase during the cue-target interval relative to the pre-cue baseline in IPS (*p* = 0.02) (IPS > EVC: *p* = 0.02, ESC > EVC: *p* = 0.03, t-tests) (**Figure 2E** left). We further visualized the spectral features of this group-level within-region CFC effect (averaged across significant electrodes in IPS) at attended locations, illustrating the peak frequencies that contributed to the most dominant coupling between alpha phase (13.4 Hz) and HFA amplitude (150 Hz) during the cue-target interval (**Figure 2C** left). At unattended locations, we also found a significant group-level increase in IPS (*p* = 7×10^−4^; alpha peak 11.4 Hz, HFA peak 170 Hz), as well as in ESC (*p* = 0.003; alpha peak 12.4 Hz, HFA peak 180 Hz) and EVC (*p* = 0.04; alpha peak 12.9 Hz, HFA peak 175 Hz) (IPS > EVC: *p* = 0.01, t-tests) (**Figure 2E** middle; **Figure 2C** middle & right). We then investigated the attentional modulation of within-region cross-frequency coupling by contrasting group-level CFC values at attended locations with those at unattended locations and found a significant attentional modulation effect in EVC (*p* = 5×10^−4^), as well as in ESC (*p* = 0.02, t-tests) (**Figure 2E** right).

To further investigate whether HFA was regulating local alpha phase or vice versa, we next used cross-frequency directionality (CFD) to measure the directionality of the coupling between alpha phase and HFA amplitude by calculating phase-slope indices (algorithms in (50), details in Methods). Group-level mean directionality was assessed through bootstrap iterations over CFD values within each area and evaluated against zero: positive mean CFD indicates that alpha phase modulates HFA power, while negative mean CFD indicates that HFA power regulates alpha phase. At attended locations, we found significantly negative group-level mean CFD in IPS (bootstrapped CFD means < 0, *p* = 1×10^−16^, t-test; median = -0.02). At unattended locations, we also found significantly negative group-level mean CFD in IPS (*p* = 4×10^−114^; median = - 0.12), EVC (*p* = 3×10^−108^; median = -0.14), and ESC (*p* = 2×10^−15^, t-tests; median = - 0.04) (**Figure 2F**; spectral features illustrated in **Figure 2D** respectively). For comparison in non-topographic areas, parietal cortex exhibited the strongest attentional modulation effect of within-region CFC, also with negative mean CFD, at unattended locations (**Figure S5**). All these negative mean within-region CFD estimates indicate that population HFA was modulating local alpha phase at the group-level. Our findings on within-region cross-frequency coupling and directionality suggest that early visual cortex and extrastriate cortex showed attentional modulation effect on within-region cross-frequency coupling, and it was HFA power envelope regulating the phase of local alpha-band activity during attentional deployment (i.e., during the cue-target interval), which might serve to attenuate sensory processing of the ignored visual space (i.e., the unattended location).

### Alpha phase modulates downstream high-frequency population activity at attended locations

After examining the neural modulation between high-frequency population activity and local alpha phase within a cortical region, we explored the effect of attentional deployment on between-region cross-frequency interactions across the visual processing hierarchy. We first assessed pair-specific CFC between alpha phase measured from one electrode and HFA amplitude measured from another electrode during the cue-target interval at contralateral attended locations (RF_desynch_) or ipsilateral unattended locations (RF_synch_), combining RFs of both electrodes for each pair with a focus on electrodes paired between visual topographic areas. Electrode pairs with an observed CFC value (sum of the maximum CFC cluster) greater than the 95% confidence interval of a shuffled surrogate distribution were considered showing significant between-region CFC effect (*p* < 0.05). For instance, between EVC and ESC at attended locations, we measured between-region CFC with alpha phase estimated from EVC electrodes and HFA amplitude from ESC electrodes (EVC_Alpha_-ESC_HFA_), as well as the reversed measure (ESC_Alpha_-EVC_HFA_), and identified the significant electrode pairs respectively (**Figure 3A**). All of these electrode pairs came from the four subjects (S1-S4) who had multiregional coverage of visual topographic areas.

We next examined the effect of attentional deployment on the group-level between-region cross-frequency coupling by comparing CFC values from significant electrode pairs during the cue-target interval to the pre-cue baseline-level CFC. At attended locations, we found a significant group-level increase for the coupling measured with alpha phase from EVC and HFA amplitude from ESC (EVC_Alpha_-ESC_HFA_: *p* = 0.03, number of electrode pairs *N* = 29; alpha peak 13.4 Hz, HFA peak 180 Hz), as well as the reversed measure (ESC_Alpha_-EVC_HFA_: *p* = 0.006, *N* = 34; alpha peak 13.9 Hz, HFA peak 165 Hz), and the coupling measured with alpha phase from ESC and HFA amplitude from IPS (ESC_Alpha_-IPS_HFA_: *p* = 0.03, *N* = 38, t-tests; alpha peak 10.4 Hz, HFA peak 180 Hz) (**Figure 3D** top, spectral features illustrated in **Figure 3B**). In contrast, at unattended locations, we found no significant group-level increase for any pair of visual topographic areas; instead, we found a significant group-level decrease for the coupling measured with alpha phase from IPS and HFA amplitude from ESC (IPS_Alpha_-ESC_HFA_: *p* = 0.009, *N* = 27; t-test) (**Figure 3D** middle). We then investigated the attentional modulation of between-region cross-frequency coupling by contrasting group-level CFC values at attended locations with those at unattended locations for each pair of visual topographic areas and found (i) between EVC and ESC, a significant attentional modulation effect for the coupling measured with alpha phase from EVC and HFA amplitude from ESC (EVC_Alpha_-ESC_HFA_: *p* = 0.003); (ii) between ESC and IPS, a significant attentional modulation effect for the coupling measured with alpha phase from ESC and HFA amplitude from IPS (ESC_Alpha_-IPS_HFA_: *p* = 0.002), as well as the reversed measure (IPS_Alpha_-ESC_HFA_: *p* = 0.04); and (iii) between EVC and IPS, a significant attentional modulation effect for the coupling measured with alpha phase from IPS and HFA amplitude from EVC (IPS_Alpha_-EVC_HFA_: *p* = 0.03, t-tests) (**Figure 3D** bottom).

To further investigate whether alpha phase in one region was modulating HFA population activity in another region or vice versa, we next estimated the directionality of between-region cross-frequency coupling by measuring the corresponding group-level between-region CFD for the identified pairs of visual topographic areas in **Figure 3B**. At attended locations, we found (i) between EVC and ESC, significantly positive group-level mean CFD for the coupling measured with alpha phase from EVC and HFA amplitude from ESC (EVC_Alpha_-ESC_HFA_: bootstrapped CFD means > 0, *p* = 0.005; median = 0.006), indicating that the phase of alpha-band activity in EVC was modulating HFA power envelope in ESC; (ii) between ESC and IPS, significantly positive group-level mean CFD for the coupling measured with alpha phase from ESC and HFA amplitude from IPS (ESC_Alpha_-IPS_HFA_: *p* = 6×10^−12^; median = 0.009), indicating that the phase of alpha-band activity in ESC was modulating HFA power envelope in IPS (for ESC_Alpha_-EVC_HFA_: bootstrapped CFD means < 0, *p* = 0.001, t-tests; median = -0.005) (**Figure 3E**, spectral features illustrated in **Figure 3C**). Our findings on between-region cross-frequency coupling and directionality suggest that alpha-band activity in lower-order visual cortex was modulating HFA power envelope in higher-order visual cortex in a phase-locking manner, potentially facilitating interareal feedforward communication during attentional deployment (i.e., during the cue-target interval) to enhance sensory processing of the attended visual space.

### Alpha rhythm contrasts interareal causal influences for target enhancement and distractor suppression

The differential local and network-level cross-frequency modulations suggest asymmetric functional connectivity during sustained visual attention in the anticipatory processing of the upcoming target versus potential distractors. To address this, we examined causal interactions in alpha-band activity and HFA responses across the visual processing hierarchy using frequency-specific Granger causality (GC) analysis to examine whether neural activity in one region was predictive of neural activity in another region, or vice versa (similar to (51)). We first identified between-region electrode pairs that showed significant GC causal influences (maximum GC cluster > 95% CI of shuffled data, *p* < 0.05) in alpha-band activity and HFA responses during the cue-target interval at attended and unattended locations for each pair of visual topographic areas (illustrated in **Figure S6** in reference to the pre-cue baseline). All of these electrode pairs came from the four subjects (S1-S4) who had multiregional coverage of visual topographic areas.

Next, we grouped significant electrode pairs for the feedforward pathway (EVC → ESC, ESC → IPS, EVC → IPS) and the feedback pathway (IPS → ESC, ESC → EVC, IPS → EVC) along the visual hierarchy, in order to investigate the effect of attentional deployment on directed frequency-specific information flow across the visual topographic cortex (illustrated in **Figure 3F**). Group-level between-region GC influence was estimated by averaging across GC values from significant electrode pairs during the cue-target interval, and then normalized to the baseline-level GC references within each pathway. At attended locations, we found a significant group-level increase during the cue-target interval relative to the pre-cue baseline for the feedforward GC influences in alpha-band activity (*p* = 0.002, t-test, number of electrode pairs *N* = 242) (**Figure 3G** left). In contrast, at unattended locations, we found a significant group-level decrease for the feedforward GC influences (*p* = 4×10^−4^, *N* = 242) and the feedback GC influences (*p* = 7×10^−6^, *N* = 236) in alpha-band activity, as well as for the feedback GC influences in HFA responses (*p* = 0.01, *N* = 240) (HFA GC feedback < feedforward: *p* = 0.003, t-tests) (**Figure 3G** middle). We then investigated the attentional modulation of between-region GC influences by contrasting group-level GC values at attended locations with those at unattended locations and found (i) in alpha-band activity, a significant attentional modulation effect for the feedforward GC influences (*p* = 3×10^−8^) and the feedback GC influences (*p* = 1×10^−4^); and (ii) in HFA responses, a significant attentional modulation effect for the feedforward GC influences (*p* = 7×10^−5^) and the feedback GC influences (*p* = 0.03) (HFA GC feedback > feedforward: *p* = 2×10^−5^, t-tests) (**Figure 3G** right). Note, the feedforward GC influences in alpha-band activity reflect increased interareal signal predictability at attended locations and decreased predictability at unattended locations. Our findings on between-region Granger causal influence suggest that, during the deployment of spatial attention, information flow mediated by alpha-band activity was enhanced along the feedforward visual pathway, facilitating functional connectivity inside the focus of attention when preparing visual cortex for the upcoming target detection; in contrast, interareal communication was suppressed with respect to the baseline outside the focus of attention, potentially attenuating the processing of distracting information.

## DISCUSSION

We used electrocorticography in human epilepsy patients to characterize neural responses during attentional deployment across the topographically organized visual system, including areas in EVC, ESC, and IPS. We observed a spatiotemporal dissociation between attention-modulated alpha desynchronization and synchronization over the visual cortex: dorsal IPS areas primarily exhibited a delayed and sustained alpha desynchronization at attended locations, while ventrolateral ESC areas primarily exhibited an earlier and sustained alpha synchronization at unattended locations. Further, we observed different neural mechanisms underlying these complementary processes: at attended locations, interareal interactions were enhanced through cross-frequency coupling along a feedforward visual pathway (EVC → ESC → IPS), possibly facilitating the processing of a spatially predictable target; in contrast, at unattended locations, intraareal interactions were enhanced through cross-frequency coupling (in EVC and ESC), and interareal interactions were suppressed, which together might serve to attenuate neural activity representing the distracting locations (schematic illustration in **Figure 3H**). Thus, our study reveals distributed and divergent neural mechanisms underlying target enhancement and distractor suppression, mediated by complex interactions between HFA and alpha-band activity during the deployment of spatial attention. The collective spatiotemporal dynamics of these alpha-mediated neural mechanisms play complementary roles in the selective processing of behaviorally relevant sensory information.

### Spatial dissociation between alpha desynchronization and synchronization

Our findings on the cortical topography of attention-modulated alpha-band activity extend previous M/EEG studies that reported alpha lateralization in parieto-occipital recording sites, characterized by contralateral decreases and ipsilateral increases in alpha power during attentional deployment (5, 12, 13, 15, 20, 23–25, 52, 53). More specifically, in previous MEG studies using the same experimental paradigm (44, 45), we reported that alpha-band activity exhibited retinotopic specificity in the visual system and exerted a Granger causal influence in the feedback direction from the frontal eye field (FEF) to primary visual cortex. Our present results, derived from ECoG data, extend beyond MEG signals by (i) revealing the distinct cortical topography of alpha power changes with precise spatial resolution, (ii) evaluating the directionality of cross-frequency interactions with an enhanced signal-to-noise ratio, and (iii) providing novel insights into the underlying neural mechanisms of interregional interactions. Here, we demonstrate that different cortical regions are linked to contralateral decreases and ipsilateral increases in alpha power, relative to the direction of attention. That is, the cortical topography of alpha desynchronization (dorsal stream) at attended locations, associated with target facilitation, is mostly distinct from that of alpha synchronization (ventrolateral stream) at unattended locations, associated with sensory suppression. This cortical dissociation between alpha desynchronization and synchronization suggests the engagement of partially overlapping, but broadly distributed, neural networks that support preparatory processing at attended and unattended locations. Previous M/EEG findings on alpha lateralization have indicated distinct neural mechanisms for target selection and distractor suppression during learning (54), proposed a selection-independent suppressive filtering mechanism supported by alpha oscillations (55), and suggested that increased alpha power reflects the suppression of distractors with increased target load (56). Our results of the distributed cortical organization and dissociated functions of alpha-band activity extend these findings by offering empirical evidence to illustrate different cortical sources for these complementary attention-selective processes.

### Temporal dissociation between alpha desynchronization and synchronization

In addition to spatial dissociation, we found a temporal dissociation between alpha contralateral desynchronization and ipsilateral synchronization. Previous EEG results have suggested an early and transient alpha power decrease during shifting of spatial attention, followed by a later and sustained alpha power increase during maintenance of attention (25). With a more precise spatial localization of cortical alpha responses, we found an early and sustained alpha power increase associated with fast and cortically distributed inhibition at unattended locations. Approximately one alpha cycle later, we observed a delayed and cortically clustered alpha power decrease at attended locations (**Figure 1E**). For inhibition, our results demonstrated local neural modulations that rapidly decoupled cortical regions possibly to block irrelevant signal transmission (**Figure 2E,3G**). For facilitation, our results demonstrated a delayed increase of functional connectivity (i.e., coupled cortical regions) which may aid relevant signal transmission (**Figure 3D,3G**). Note, the reestablished and maintained alpha desynchronization during target selection (**Figure 1E**) possibly reflected the event-related potential (ERP) contribution power in the alpha band. Our results provide empirical evidence to support the critical involvement of alpha lateralization in both facilitatory and inhibitory processes, further demonstrating that these complementary processes are not only cortically distributed but also temporally dissociated.

### Local cross-frequency modulation for distractor filtering

We next demonstrate different local and interareal interactions associated with neural processing at attended and unattended locations. To characterize local neural modulation at unattended locations, our study advances previous work on the spatial tuning and topographic properties of alpha-band activity (45, 47) and high-frequency broadband activity (46), specifically examining modulations of their cross-frequency interactions during the deployment of spatial attention (as proposed by the alpha pulsed-inhibition hypothesis in (8)). We found that the power envelope of HFA drives the alpha phase of local circuits, particularly at unattended locations associated with distractor inhibition (**Figure 2E,2F**). This is consistent with previous human ECoG results showing that cortical high-gamma activity regulates local alpha phase within posterior-ventrolateral occipital/temporal cortical sites during resting state (50). Our data offers novel empirical evidence to support the notion that such coordinated firing patterns of neuronal populations within a cortical region, as reflected in HFA, may play a critical role in the emergence or amplification of local oscillatory activity as a function of attentional deployment. However, these results challenge the conventional view that slow oscillations modulate the excitability of local neuronal populations (57). Furthermore, the interareal interactions as demonstrated in Granger causal influences support the notion of disrupted information transfer across the visual hierarchy to attenuate distractor processing. At unattended locations, although alpha power increases locally, interareal cortical interactions (i.e., functional connectivity) in the alpha-band decrease (**Figure 3G**). This suggests that the fast and local modulation of alpha ipsilateral activity may act to quickly decouple irrelevant cortical regions, thereby suppressing signal transmission associated with the representation of potential distractor locations. Our findings on the early local processes of distant distractor filtering (suppressing opposite locations) further extend previous literature showing an early-stage filtering of adjacent distractors (suppressing neighboring objects) at the level of the receptive field as proposed by the biased competition theory of selective attention (58–60).

### Network cross-frequency modulation for target enhancement

At attended locations, we found that alpha phase in EVC modulates the high-frequency power envelope in ESC, and that alpha phase in ESC further modulates HFA in IPS (**Figure 3D,3E**). This directed, interareal cross-frequency interaction provides empirical evidence to support previous hypotheses on alpha rhythmic modulation of downstream population neuronal firing (8), by revealing an underlying network modulation associated with facilitated sensory processing. Note, when considering HFA as a proxy for population spiking, the alpha-HFA phase-amplitude coupling demonstrates a comparable pattern to the spike-LFP phase coupling observed in single/multi-unit activity, suggesting that population spiking activity is modulated as a function of alpha phase. The Granger causal influences data further support enhanced information flow in the alpha-band along the feedforward visual pathway to facilitate upcoming target processing (**Figure 3G**). This is in line with recent findings that relate feedforward alpha propagation to ongoing visual processing (61), challenging previous work showing that alpha/beta rhythms predominantly exert feedback influences on visual cortex in target detection (62, 63). We provide an alternative perspective on how information transmission over the visual hierarchy is enhanced through increased feedforward connectivity in the alpha band, potentially elevating the cortical interconnection of relevant regions representing the attended space. Further, recent ECoG findings in marmoset monkeys have demonstrated a traveling wave of theta/alpha oscillations from posterior parietal to ventral extrastriate cortex via primary visual cortex during free viewing (64). Our present results, demonstrating a ventrolateral alpha feedforward flow (EVC → ESC → IPS), in conjunction with our previous findings of a dorsal alpha feedback signal (FEF → EVC in (44)), suggest a flow of visual processing involving the alpha-band activity between the occipital visual system and the frontoparietal attention network, utilizing both bottom-up and top-down processes. Together, our cross-frequency analyses of within-region and between-region coupling reveal differential local and network neural mechanisms of attentional modulation (65), illustrating the effective gating of information flow across the visual processing network through alpha-mediated facilitatory or inhibitory control.

In summary, alpha-band activity mediates the selective gating of visual processing, utilizing enhanced functional connectivity through interareal neural modulation at attended locations and attenuating distractor processing with suppressed network connectivity through local neural modulation at unattended locations. Our study reveals both facilitatory and inhibitory roles of cortically distributed alpha-band activity during the deployment of visual spatial attention related to target enhancement at attended locations and distractor suppression at unattended locations.

## SUPPLEMENTARY INFORMATION

### METHODS

#### Subjects

Intracranial recordings were obtained from 8 human subjects (S1-S8, 6 males; age: mean ± SEM = 35 ± 5 years), who underwent pre-surgical epilepsy evaluation at Stanford University Medical Center, Johns Hopkins Hospital, UC Irvine Medical Center, and Oakland Children’s Hospital (demographic details are available in (1)). All subjects provided written informed consent to participate in this study, which was approved by the Institutional Review Boards of the participating institutions. Subjects had been discontinued of anti-epileptic medications and were seizure free for at least 5 hours before testing. All subjects had normal or corrected-to-normal vision.

#### Electrophysiological recordings

Subjects were implanted intracranially with grid and/or strip subdural platinum electrodes, covering extensive parts of occipital, parietal, frontal, and temporal cortex over the left (7 subjects) and right (1 subject) cerebral hemispheres. The majority of implanted electrodes were 4 mm in diameter embedded in a flexible silicon sheet, with an exposed recording diameter of 2 mm and an inter-electrode spacing of 10 mm. All implantations were guided entirely by clinical requirements and considerations.

Electrophysiological recordings were performed using a Tucker-Davis Technologies (TDT) recording system at Stanford, a Stellate Harmonic or a Blackrock recording system at Johns Hopkins, and Nihon Kohden (NK) recording systems both at UC Irvine and Oakland. Signals were sampled at 3052 Hz (TDT), 1000 Hz (Stellate), 10000 Hz (Blackrock), or 5000 Hz (NK), amplified and bandpass filtered (0.5-300 Hz at Stanford; 0.1-350 Hz (Stellate) or 0.3-2500 Hz (Blackrock) at Johns Hopkins), using a scalp ground and a subdural reference electrode, selected as the most electrographically silent channel outside of the clinically identified seizure zone (recording details at each institution are available in (1)).

#### Electrode localization

We reconstructed electrode locations by first aligning a post-operative CT scan image of the implanted electrodes with a pre-operative high-resolution structural MRI using SPM8 (2). We then localized three-dimensional coordinates of each electrode in the co-registered CT image and further corrected for minor cortical shifts following surgery (similar to (3)). Electrode locations were mapped onto a rendering of cortical surfaces reconstructed from each subject’s structural MRI using FreeSurfer (4, 5) and further converted to a standard surface template using SUMA (6) and AFNI (7).

For anatomical localization of electrodes within the visual system, we used a probabilistic atlas of visuospatial topographic areas based on fMRI retinotopic mapping data (8). Using the maximum probability map, each electrode that overlapped with the atlas was assigned to the topographic area with the highest probability. Electrodes outside the visuospatial topographic areas were localized using the Harvard-Oxford cortical parcellation based on anatomical markers (9). Coverage of individual patients is depicted on a standard brain surface in **Figure S1**.

#### Experimental task and behavioral analysis

Subjects performed a variant of an Eriksen flanker task (1, 10–12). Visual stimuli were presented on a laptop computer placed approximately 80 cm away from the subject’s eyes using Presentation software (Neurobehavioral Systems, Inc.). Each trial started with the presentation of a central fixation point for 1100 ms, followed by a circular spatial cue briefly displayed for 100 ms in a pseudo-randomly selected location arranged in a circular manner around the fixation point at a fixed eccentricity of 7°. After a variable delay interval of 300-700 ms, a circular array of equally spaced barrel and bowtie shapes was presented for 2000 ms or until a response was recorded. Subjects were instructed to respond with a left or right button press to indicate whether a barrel or bowtie target shape was presented at the cued location. Either congruent or incongruent shapes could flank the target shape. A signaling tone was delivered upon completion of each trial to indicate a correct or incorrect response. Trials were separated by a 2 s inter-trial interval. After a training block to familiarize themselves with the task, subjects performed 3-6 blocks in total, each containing 50 trials, for the actual experiment. The number of cue and target locations was 8, 14, or 16 for different subjects to adjust for task difficulty (experimental details of each subject are available in (1)). Subjects were instructed to maintain central fixation throughout the entire trial, and at some recording sites their eye movements were visually monitored by the experimenter with the assistance of video recordings in the patient room. No systematic saccadic eye movements were observed during the experiment.

For each subject, we computed behavioral accuracy (i.e., the proportion of correct trials relative to the number of all trials) and mean reaction times (RTs) averaged across all correct trials (details of behavioral performance for each subject are available in (1)). We further analyzed the behavioral accuracy as a function of the flanker condition (congruent vs. incongruent flanker stimuli) to confirm that subjects were performing the task as instructed. For the subsequent analyses of neural data, only trials with correct responses with RTs ≤ 3 SD from the mean (within subject) were included.

#### ECoG data preprocessing

All analyses were performed offline using EEGLAB (13), FieldTrip (14), and customized scripts written in MATLAB (MATLAB R2020a, The Mathworks Inc., Natick MA). Prior to data processing, electrodes clinically identified within the ictogenic zone and those observed as electrically corrupted during recording were eliminated from subsequent data analyses (median = 1.3% of total electrodes, min = 0%, max = 10.2%). Electrodes were further excluded from common average reference whose overall voltage variance was greater than five times or less than one-fifth of the median variance across all electrodes (median = 5.7% of total electrodes, min = 0.9%, max = 32.3%), those with more than one large spike jump (> 80 µV difference between adjacent time points) per second on average (median = 0.4% of total electrodes, min = 0%, max = 9.4%), and those whose power spectrum did not show the typical 1/f power spectrum based on visual inspection (median = 0.8% of total electrodes, min = 0%, max = 7.8%). All non-pathological electrodes were then notch filtered at 60 Hz and its harmonics (120 Hz and 180 Hz) to remove electric interference, then re-referenced to the common average of the filtered signals of the non-excluded electrodes. The re-referenced signal at each electrode was decomposed into 55 bandwidths by filtering the signal with 55 custom-made band-pass filters; each filter had a central frequency chosen from 1 to 250 Hz range with equal log-distance spacing (similar to (15)). We applied the Hilbert transform to each filtered time series and acquired the instantaneous amplitude. The amplitude of each band signal was normalized by its own mean, then these normalized amplitude time series were averaged together, yielding a single amplitude time course for alpha-band activity (8-20 Hz) and broadband high-frequency activity (HFA, 70-200 Hz). Note that this approach accounts for the 1/f signal drop off in HFA power at increasing frequencies.

#### Recording sites exhibiting task-related alpha responses

Our study probed the effects of attentional modulation during the cue-target delay, i.e., when spatial attention was allocated in anticipation of target appearance and during a period of pure cognitive state without a change in the sensory environment. We thus focused our analyses on modulation of alpha-band activity during a 300 ms time window of the cue-target interval (i.e., cut-off of 8 Hz). Alpha power during the cue-target delay (300 ms before target onset) was first z-score normalized relative to the pre-cue baseline (300 ms before cue onset) of each trial, at each of the peripherally cued locations and for each electrode. Trials with outlier delay-related alpha power (i.e., with outlier time points > mean ± 6 SD) were eliminated (median = 10.6% of previously selected trials, min = 2.2%, max = 22.4%). We then averaged alpha power during the cue-target delay and the pre-cue baseline within each trial, yielding one mean alpha value for delay and baseline per trial. Based on previous EEG studies on attentional modulation of alpha power that showed spatially-specific decreases at attended locations and increases at unattended locations (16–18), we defined an alpha desynchronization ECoG response field (RF_desynch_) by identifying visual field locations that showed the greatest alpha power decrease when attention was deployed within the contralateral visual field. Similarly, we defined an alpha synchronization response field (RF_synch_) by identifying visual field locations that showed the greatest alpha power increase when attention was deployed within the ipsilateral visual field. Specifically, as the number of cue locations varied from 8 to 16 for different subjects, alpha delay power was averaged across 2-3 neighboring locations across subjects, covering approximately 45-51 degrees of visual field (i.e., the size of alpha response fields). The visual field locations that showed the greatest alpha power decrease or increase, as compared to other combinations of neighboring locations in the same hemifield, were identified as alpha RF_desynch_ or RF_synch_ respectively. Vertical top and bottom locations were not included in the selection of response field locations in order to avoid double-counting the same set of trials for the attended and unattended conditions. Note that, within each subject, the size of alpha response fields was kept the same for each of the cortical areas, so that peak alpha responses across different brain areas could be compared in an unbiased way. We then performed non-parametric permutation tests to identify electrodes that showed significant delay-related changes in alpha-band activity. These tests were run separately for trials pooled over the identified alpha RF_desynch_ and RF_synch_ to test for differences in alpha power between the delay group and the baseline group, randomly reassigning the group labels 1000 times. One-sided *p*-values were calculated based on distributions of values generated from the permutation tests; electrodes with significant alpha power decreases at RF_desynch_ or increases at RF_synch_ (*p* < 0.05) were considered as task-responsive cortical sites that showed delay-related alpha desynchronization or synchronization relative to pre-cue baseline, respectively.

#### Identifying electrodes with attention-modulated spatial selectivity

To further investigate whether electrodes that showed delay-related alpha responses also exhibited spatially selective attention-modulated changes in alpha power, we quantified whether the distributions of mean alpha power at RF_desynch_ or RF_synch_ were different from those at the corresponding opposite visual field locations by calculating alpha selectivity indices (SI). Specifically, alpha RF_desynch_ SI values measured the distance between the means of alpha power at RF_desynch_ and the corresponding opposite visual field locations, normalized in units of the pooled standard deviation of both distributions; alpha RF_synch_ SI values were calculated similarly using trials cued at RF_synch_ and the corresponding opposite locations. Electrodes were sorted by their alpha RF_desynch_ and RF_synch_ SI values and visualized in a two-dimensional space for each cortical area. To identify electrodes that showed attention-modulated spatially selective changes in alpha power, we first estimated the SI significance threshold by bootstrapping SI values of the non-responsive electrodes (i.e., electrodes that neither showed significant alpha power decreases at RF_desynch_ nor increases at RF_synch_ during the cue-target delay) 200 times within each cortical area. From the pool of electrodes that showed significant delay-related changes in alpha power, we identified those with alpha RF_desynch_ SI values less than the one-sided 95% confidence interval of the estimated SI distribution (bootstrap mean - 1.645 × bootstrap SD) as showing significant spatially selective alpha desynchronization at attended locations, and those with alpha RF_synch_ SI values greater than the one-sided 95% confidence interval (bootstrap mean + 1.645 × bootstrap SD) as showing significant spatially selective alpha synchronization at unattended locations.

We next visualized the cortical distribution of attention-modulated alpha desynchronization and synchronization using electrodes that showed spatially selective attention-modulated changes in alpha power during the cue-target delay. Specifically, we rendered the alpha RF_desynch_ and RF_synch_ SI values from all subjects onto a standard brain surface to visualize the cortical topography of attention-modulated spatially selective alpha desynchronization and synchronization. The topography was spread cortically using a Gaussian kernel of 4 cm. We further investigated the temporal dynamics and sustainability of attention-modulated alpha cortical topography by computing alpha SI values within three consecutive 100 ms time windows (approximately 1 cycle of alpha oscillations at 10 Hz) during the cue period (300 ms after cue onset) and during the delay period (300 ms before target onset) from the identified ‘Selective’ electrodes. We also rendered these alpha SI values from all subjects onto a standard brain surface within each 100 ms time windows to illustrate the temporal dynamics of alpha cortical topography after the presentation of the spatial cue and during the maintenance of spatial attention.

Lastly, we visualized the time course of attention-modulated alpha desynchronization and synchronization throughout the entire task trial using electrodes that showed spatially selective changes in alpha delay activity. We first acquired mean alpha power time series at each electrode within each task epoch by averaging across trials, then smoothed the averaged time series with a linear interpolation using the least common multiple of the subjects’ sampling rates to align across all electrodes and further downsampled to 1000 Hz. We performed t-tests on the aligned alpha power time series across electrodes at each time point to identify time periods when alpha power at RF_desynch_ or RF_synch_ were both significantly different than zero (i.e., baseline-level alpha power) and significantly different than those at the corresponding opposite visual field locations (*p* < 0.05, Bonferroni corrected for multiple comparisons).

#### Identifying electrodes with cross-frequency phase-amplitude coupling

Broadband high-frequency activity (HFA, 70-200 Hz) has been shown to be highly correlated with population neuronal spiking (19–21) or superficial dendritic processes (22, 23). In order to investigate the relationship between HFA and alpha responses, we examined whether the HFA power envelope (a proxy for population spiking) modulated the phase of alpha-band activity or vice versa by first measuring cross-frequency coupling (CFC), both within cortical regions and between cortical regions. For each electrode (within-region) or pair of electrodes (between-region), we used trials cued at alpha RF_desynch_ or RF_synch_ (combining RFs of both electrodes for between-region measurement) and segmented them to isolate the cue-target delay interval, as well as the pre-cue baseline for reference. We z-score normalized the segmented signals to the pre-cue baseline and further excluded trials with outlier voltage to increase the signal-to-noise ratio (median = 0.01% of previously selected trials, min = 0%, max = 0.03%). We then decomposed the normalized signals into different frequency components using a fast Fourier transform (FFT), examining a broad frequency range from 1 to 30 Hz in steps of 0.5 Hz for phase estimates and from 30 to 200 Hz in steps of 5 Hz for power measurements. The strength of CFC was calculated by computing the coherence between low-frequency phase and high-frequency power envelope for paired frequency bins (algorithms adapted from (24)). In order to quantify the dominant coupling effect with consideration of spectral variance across electrodes or pairs of electrodes, we set an electrode-specific or pair-specific threshold to identify the maximum cluster of CFC estimates exceeding the threshold, such that the CFC maximum cluster was above min_CFC_ + (max_CFC_ - min_CFC_) × 0.8. We then calculated the sum of the maximum cluster and referred to as CFC value for subsequent analyses. To identify electrodes or pairs of electrodes with significant cross-frequency coupling, we generated a surrogate distribution by computing CFC values after first randomly shuffling the alpha phase estimates relative to HFA amplitude across trials for 1000 times, using shuffle-specific threshold. Electrodes or pairs of electrodes with an observed CFC value greater than the 95% confidence interval of the shuffled distribution were considered statistically significant (*p* < 0.05). Note, for between-region CFC analysis, we paired electrodes between visual topographic areas in subjects with simultaneous coverage; each pair of electrodes was tested twice with one electrode serving as the seed to estimate alpha phase while the other for HFA amplitude, and vice versa.

To quantify group-level cross-frequency coupling, we averaged across CFC values from significant electrodes or pairs of electrodes for within-region or between-region CFC, after excluding outliers > 3 SD from the mean (mean & SD were estimated through 200 bootstrap iterations over CFC values within each area or pair of areas, accounting for the difference in numbers of data points). For within-region CFC analysis, we further weighted the group-level CFC by the percentage of significant electrodes within each area to better illustrate the magnitude of group-level within-region CFC effect. We next investigated the effect of attentional deployment on cross-frequency coupling by comparing group-level CFC values during the cue-target delay to those during the pre-cue baseline for each area or pair of areas. To assess the group-level statistical significance, we performed t-tests on the normalized CFC values to evaluate whether the group-level within-region or between-region CFC was either different than zero (i.e., baseline-level CFC) or different from other areas or pairs of areas. The effect of attentional modulation on within-region or between-region CFC was further examined by contrasting group-level CFC values at attended locations (alpha RF_desynch_) with those at unattended locations (alpha RF_synch_) within or between cortical areas.

#### Measuring directionality of neuronal modulation and functional connectivity

To further investigate whether the HFA power envelope modulated the phase of alpha-band activity or vice versa, we next used cross-frequency directionality (CFD) to measure the directionality of the coupling between alpha phase and HFA amplitude, both within cortical regions and between cortical regions. For each electrode or pair of electrodes that showed significant CFC effect, we calculated phase-slope indices between low-frequency phase and high-frequency amplitude; frequency bandwidth to estimate phase slope was set to 2 Hz at central phase frequency from 8 to 20 Hz in 1 Hz steps. This approach allowed us to infer whether one frequency component (i.e., alpha phase or HFA amplitude) was leading or lagging the other based on the assumption that a constant time lag could be translated into phase differences (24, 25). To quantify the CFD value of the dominant coupling effect within each electrode or pair of electrodes, we applied a spectral filter of the maximum CFC cluster (see previous section) and calculated the sum of the filtered CFD estimates. We then assessed the group-level mean directionality through 200 bootstrap iterations over CFD values within each area or pair of areas, and further evaluated the distribution of bootstrapped CFD means against zero by performing t-tests. Positive mean CFD indicates that alpha phase modulates HFA power, while negative mean CFD indicates that HFA power regulates alpha phase, either within a cortical region or from one region to another.

To further explore the causal influence of information flow along the visual hierarchy during sustained attention, we applied frequency-specific Granger causality (GC) analysis to estimate causal interactions in alpha-band activity and HFA responses in subjects with multiregional coverage of visual topographic areas. For each pair of electrodes, we used trials cued at alpha RF_desynch_ or RF_synch_ (combining RFs of both electrodes) and segmented them to the cue-target delay and the pre-cue baseline intervals, then z-score normalized the segmented signals to the pre-cue baseline and further excluded trials with outlier voltage. To calculate spectrally resolved GC influences, we fit multivariate autoregressive models to the trial data using the BSMART toolbox (26), with a model order of 8 for the computation of autoregressive coefficients, and computed GC estimates within the alpha-band frequencies (8-20 Hz) and HFA frequencies (70-200 Hz) in 1 Hz steps. In order to quantify the dominant GC influences with consideration of spectral variance across electrode pairs, we set a pair-specific threshold to identify the maximum cluster of GC estimates exceeding the threshold, such that the GC maximum cluster was above min_GC_ + (max_GC_ - min_GC_) × 0.8. We then calculated the sum of the maximum cluster and referred to as GC value for subsequent analyses. To identify pairs of electrodes with significant GC influences, we generated a surrogate distribution by computing GC values after first randomly shuffling the trial labels for 1000 times, using shuffle-specific threshold. Electrode pairs with an observed GC value greater than the 95% confidence interval of the shuffled distribution were considered statistically significant (*p* < 0.05). To quantify group-level between-region GC influences, we averaged across GC values from significant electrode pairs for the feedforward pathway (EVC → ESC, ESC → IPS, EVC → IPS) and the feedback pathway (IPS → ESC, ESC → EVC, IPS → EVC) along the visual hierarchy, after excluding outliers > 3 SD from the mean (mean & SD were estimated through 200 bootstrap iterations over GC values within each visual pathway, accounting for the difference in numbers of data points). Outlier electrode pairs were also excluded with outlier GC estimates at any frequency bin across electrode pairs. We then investigated the effect of attentional deployment on GC influences by comparing the group-level GC values during the cue-target delay to those during the pre-cue baseline, for alpha-band activity and HFA responses in each visual pathway. To assess the group-level statistical significance of GC influences, we performed t-tests on the normalized GC values to evaluate whether the group-level feedforward or feedback GC influence was either different than zero (i.e., baseline-level GC) or different from each other. The effect of attentional modulation on between-region GC influences was further examined by contrasting group-level GC values at attended locations (alpha RF_desynch_) with those at unattended locations (alpha RF_synch_) for each visual pathway.

---

**Figure S1.**
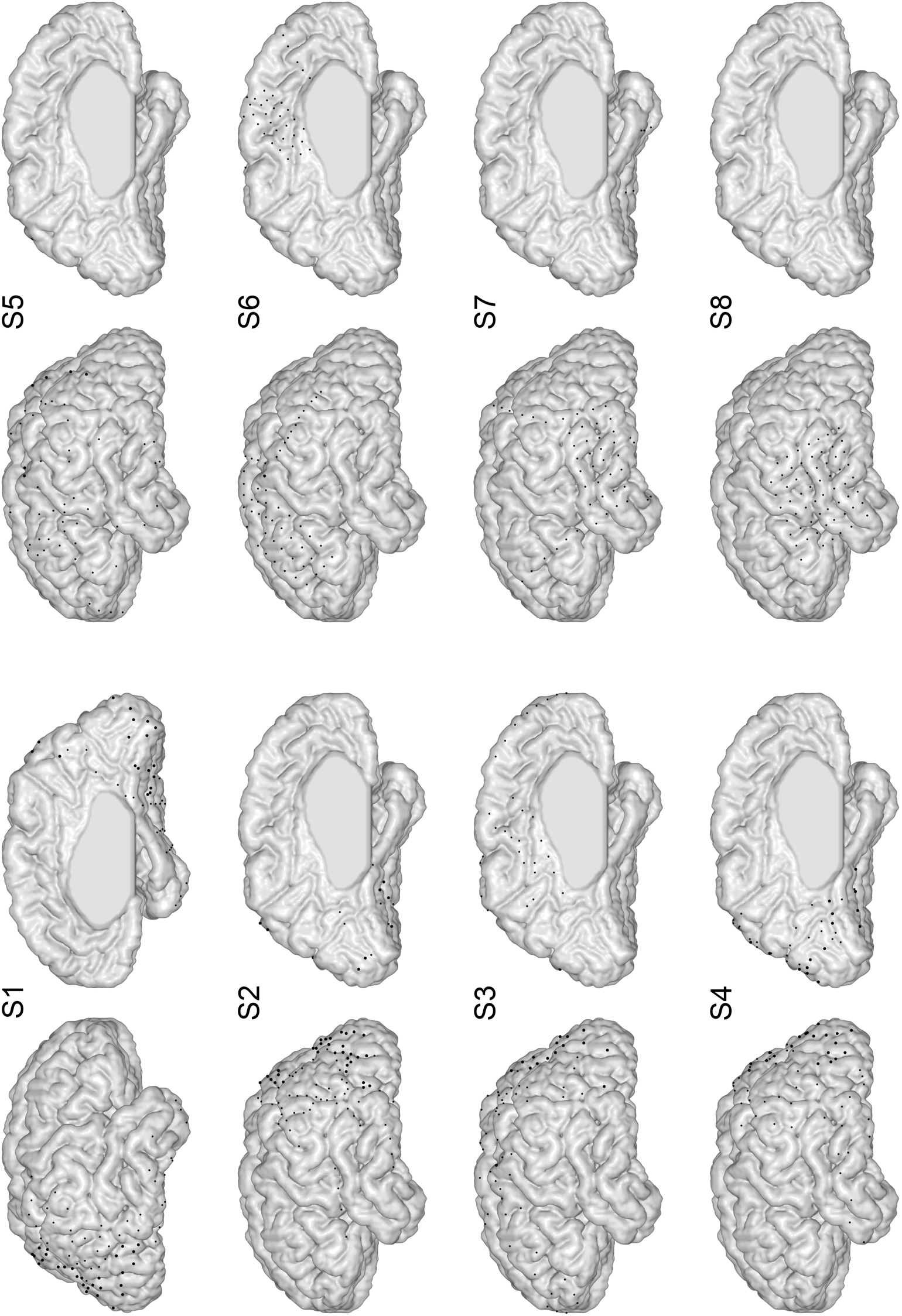
Electrode coverage of individual patients. Electrode coverages of individual patients are depicted on a standard brain surface shown from the lateral and medial views for subject S1 with right hemisphere implantations and subjects S2-S8 with left hemisphere implantations. Large dots denote electrodes located in visual topographic cortex, and small dots those located in non-topographic cortex.

**Figure S2.**
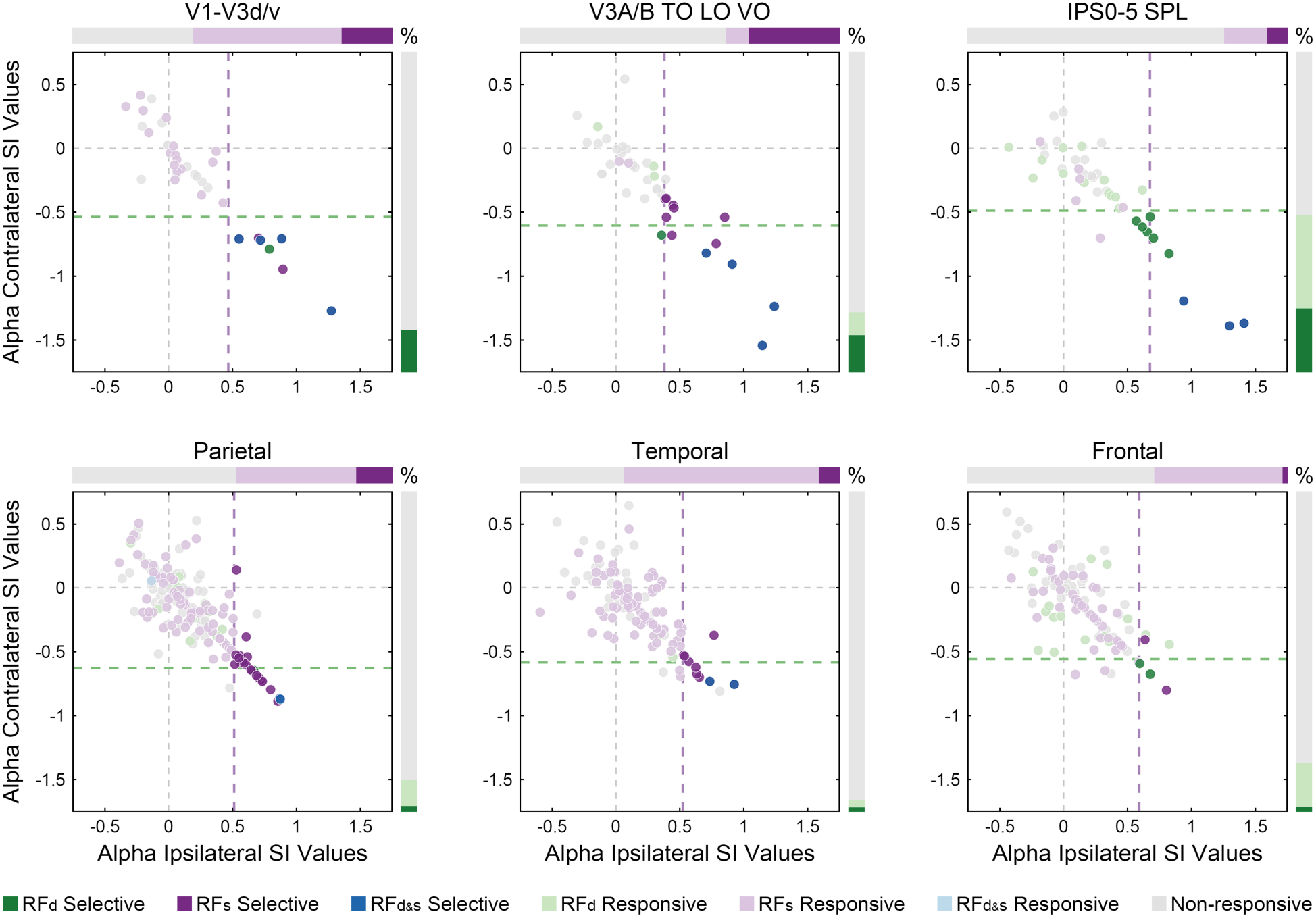
Spatial selectivity of attention-modulated alpha responses. Electrodes sorted by alpha RF_desynch_ and RF_synch_ SI values during the cue-target delay, illustrated for each of the visual topographic and non-topographic areas. For each panel, the contralateral-axis shows alpha RF_desynch_ SI values with the green dashed line indicating the alpha RF_desynch_ SI threshold for significant spatial selectivity (estimated by bootstrapping SI values of the ‘Non-responsive’ electrodes), while the ipsilateral-axis shows alpha RF_synch_ SI values with the purple dashed line indicating the alpha RF_synch_ SI threshold. Electrodes are color-coded based on their alpha-band activity during the cue-target delay: electrodes that did not show any significant delay-related changes in alpha-band activity (‘Non-responsive’) are marked in gray; electrodes that showed significant delay-related changes in alpha-band activity (‘Responsive’) are marked in light green for the attended condition associated with alpha desynchronization (‘RF_d_’), in light purple for the unattended condition associated with alpha synchronization (‘RF_s_’), and in light blue for both conditions (‘RF_d&s’_); electrodes that showed significant spatially selective changes in alpha-band activity (‘Selective’) are marked in dark green for the attended condition, in dark purple for the unattended condition, and in dark blue for both conditions. The right-side vertical bars and the upside horizontal bars illustrate the percentage of alpha ‘Non-responsive’, ‘Responsive’, and ‘Selective’ electrodes within each cortical area for the attended and unattended conditions respectively. Note that ‘RF_d&s_’ electrodes are included in ‘RF_d_’ and ‘RF_s_’ electrodes for the illustration of electrode percentage. Numbers of electrodes and contributing subjects for each category are listed in **Table S1**.

**Figure S3.**
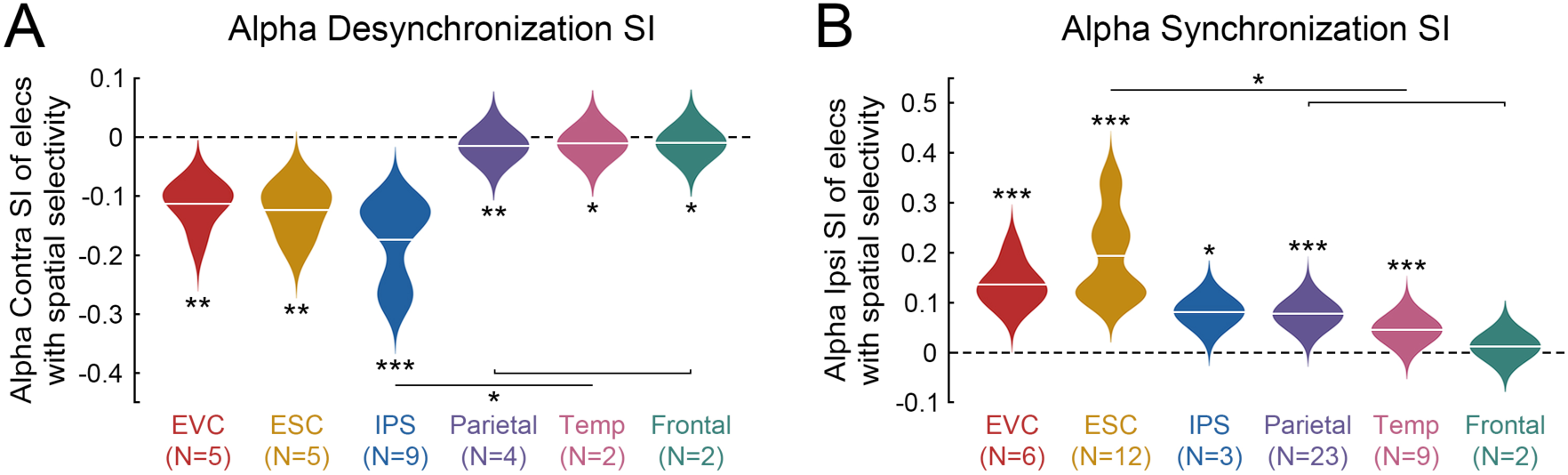
Strengths and dissociation of attention-modulated alpha responses. **A:** Group-level alpha RF_desynch_ SI values from electrodes that showed spatially selective alpha responses during the cue-target interval at attended locations, weighted by the percentage of significant electrodes within each topographic and non-topographic areas, where alpha RF_desynch_ SI were found showing the strongest contralateral desynchronization in IPS (IPS < 0: p = 7×10^−5^; EVC < 0: p = 0.002, ESC < 0: p = 0.003, parietal < 0: p = 0.001, temporal < 0: p = 0.01, frontal < 0: p = 0.04), significantly less than non-topographic cortex (IPS < parietal: p = 1×10^−3^, IPS < temporal: p = 0.01, IPS < frontal: p = 0.01). **B:** At unattended locations, alpha RF_synch_ SI were found showing the strongest ipsilateral synchronization in ESC (ESC > 0: p = 1×10^−5^; EVC > 0: p = 4×10^−4^, IPS > 0: p = 0.01, parietal > 0: p = 1×10^−18^, temporal > 0: p = 3×10^−7^), significantly greater than non-topographic cortex (ESC > parietal: p = 4×10^−7^, ESC > temporal: p = 8×10^−5^, ESC > frontal: p = 0.02). All statistical significance was established using t-tests (* p < 0.05, ** p < 0.01, *** p < 0.001).

**Figure S4.**
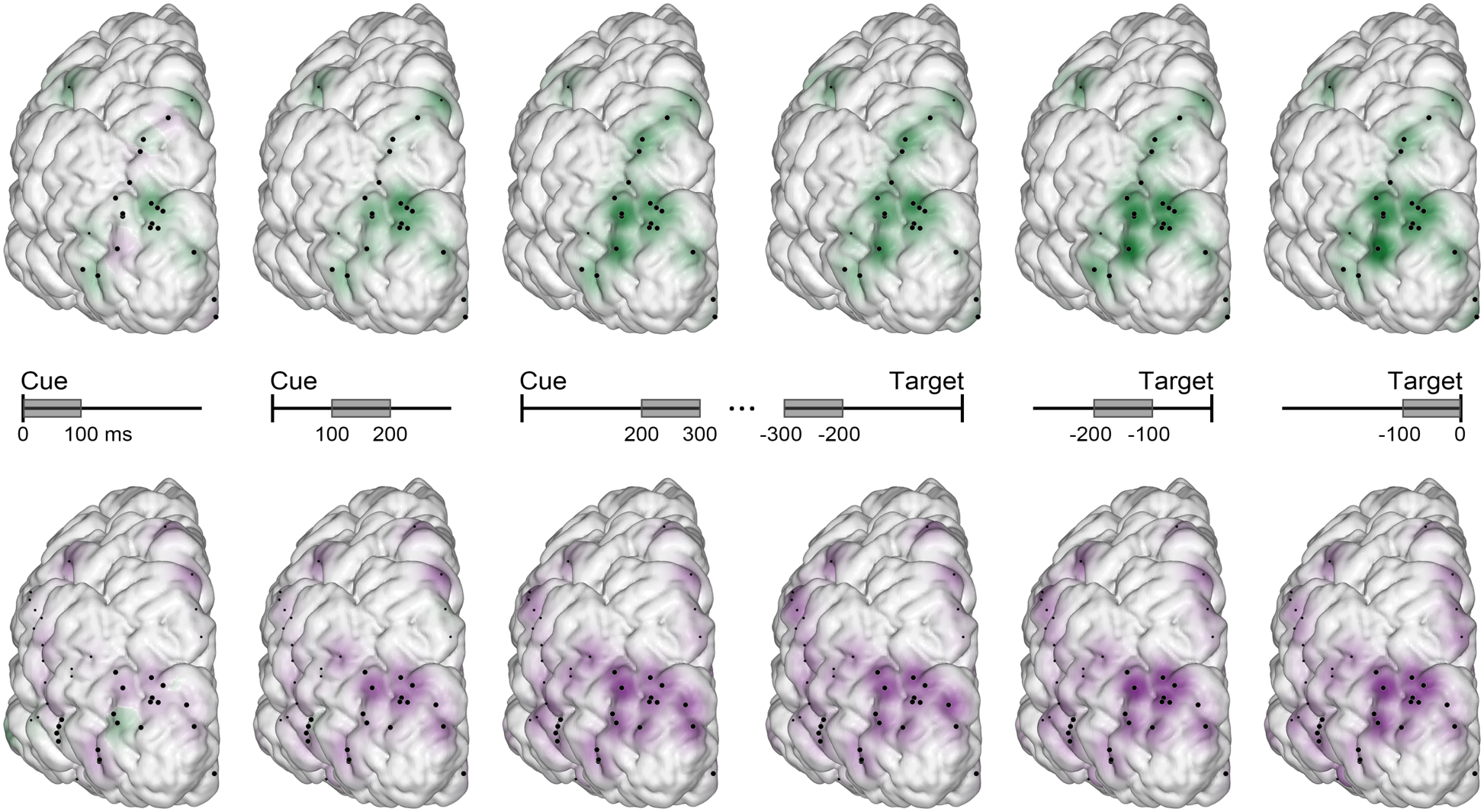
Temporal dynamics of attention-modulated alpha topography. Cortical topography of alpha RF_desynch_ and RF_synch_ SI values from electrodes that showed spatially selective changes in alpha-band activity during the cue-target delay, computed within three consecutive 100 ms time windows during the cue period aligned to cue onset (left panels) and during the delay period aligned to target onset (right panels), rendered across all subjects onto a standard brain surface shown from the posterior view. Attention-modulated alpha desynchronization is illustrated in green showing a build-up of and sustained dorsal distribution (upper panels), whereas attention-modulated alpha synchronization is illustrated in purple showing a build-up of and sustained ventrolateral distribution (lower panels). Black dots denote cortical localization of electrodes that showed spatially selective changes in alpha-band activity; large dots denote electrodes located in visual topographic cortex, and small dots for electrodes located in non-topographic cortex.

**Figure S5.**
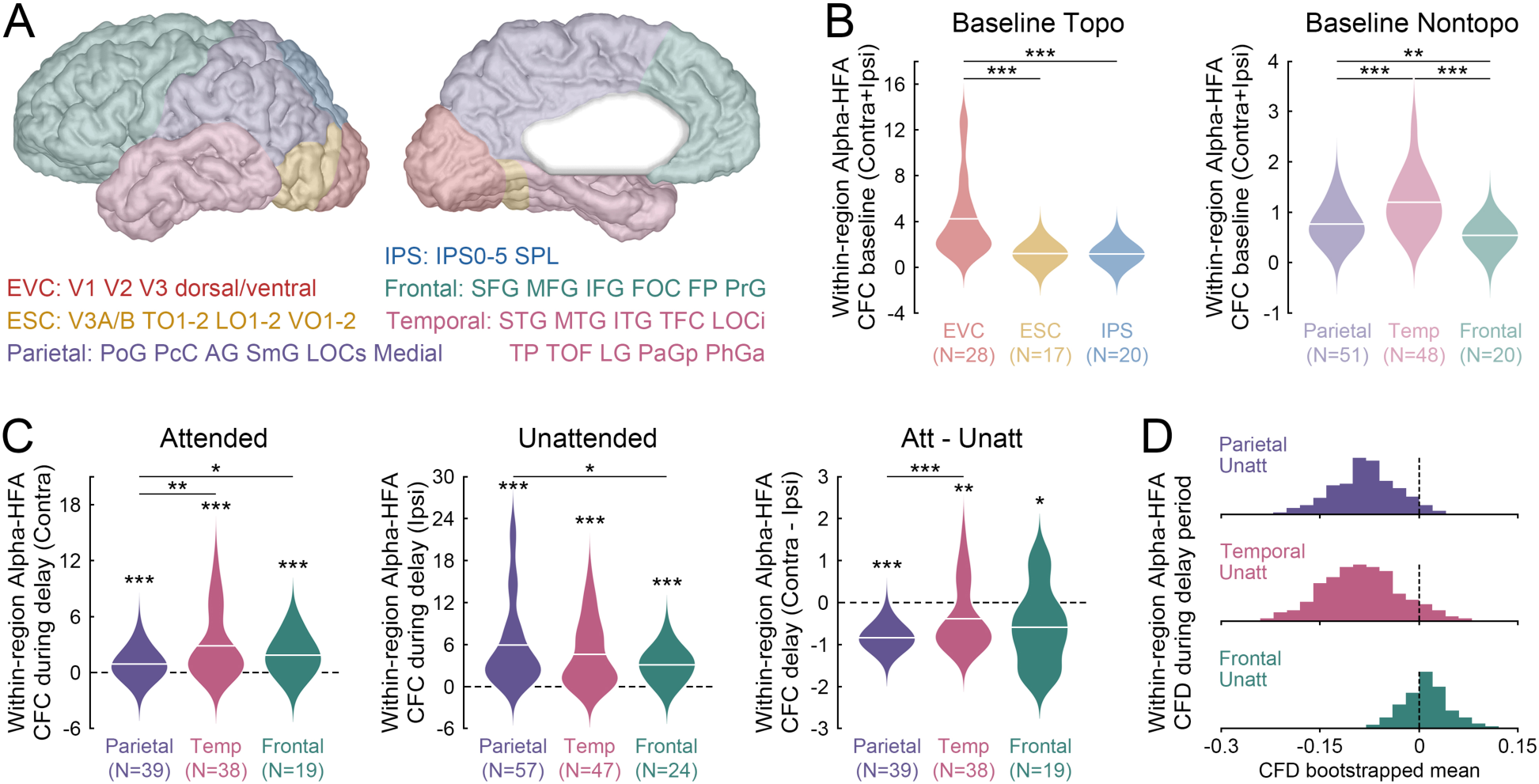
Attentional modulation of cross-frequency interactions within cortical regions in non-topographic areas. **A:** Color-coded ROIs are listed for each of the visual topographic areas identified using our probabilistic atlas of the human visual system, and for each of the non-topographic areas labelled using the Harvard-Oxford cortical parcellation. **B:** Group-level within-region CFC during the pre-cue baseline for each visual topographic areas, with the strongest baseline-level CFC found in EVC (EVC > ESC: p = 7×10^−4^, EVC > IPS: p = 2×10^−4^) (left panel). In non-topographic areas, the strongest baseline-level CFC was found in temporal cortex (temporal > parietal: p = 9×10^−6^, temporal > frontal: p = 4×10^−6^, parietal > frontal: p = 0.006) (right panel) (numbers of contributing electrodes enclosed in parentheses, combining baselines at attended and unattended locations). **C:** Group-level within-region CFC during the cue-target delay at attended locations for each non-topographic areas, where significant group-level increases were found in parietal cortex (p = 5×10^−5^), temporal cortex (p = 1×10^−5^), and frontal cortex (p = 5×10^−4^); temporal > parietal (p = 0.002), frontal > parietal (p = 0.02) (left panel). At unattended locations, significant group-level increases of within-region CFC were also found in parietal cortex (p = 8×10^−10^), temporal cortex (p = 1×10^−8^), and frontal cortex (p = 2×10^−7^); parietal > frontal (p = 0.03) (middle panel). The attentional modulation effect of within-region CFC was estimated by contrasting group-level CFC values at attended locations with those at unattended locations, where significant attentional modulation effects were found in parietal cortex (p = 8×10^−26^), temporal cortex (p = 0.004), and frontal cortex (p = 0.01); |parietal| > |temporal| (p = 7×10^−4^) (right panel). **D:** Group-level mean directionality of within-region CFC was estimated through bootstrap iterations over CFD values from significant electrodes within each area at unattended locations. The distributions of the bootstrapped CFD means are illustrated and evaluated against zero, where significantly negative group-level mean CFD was found in parietal cortex (bootstrapped CFD means < 0, p = 2×10^−62^; median = -0.08) and temporal cortex (p = 4×10^−53^; median = -0.09) indicating that HFA power regulates alpha phase, while significantly positive group-level mean CFD was found in frontal cortex (p = 1×10^−5^; median = 0.01) indicating that alpha phase modulates HFA power. All statistical significance was established using t-tests (* p < 0.05, ** p < 0.01, *** p < 0.001).

**Figure S6.**
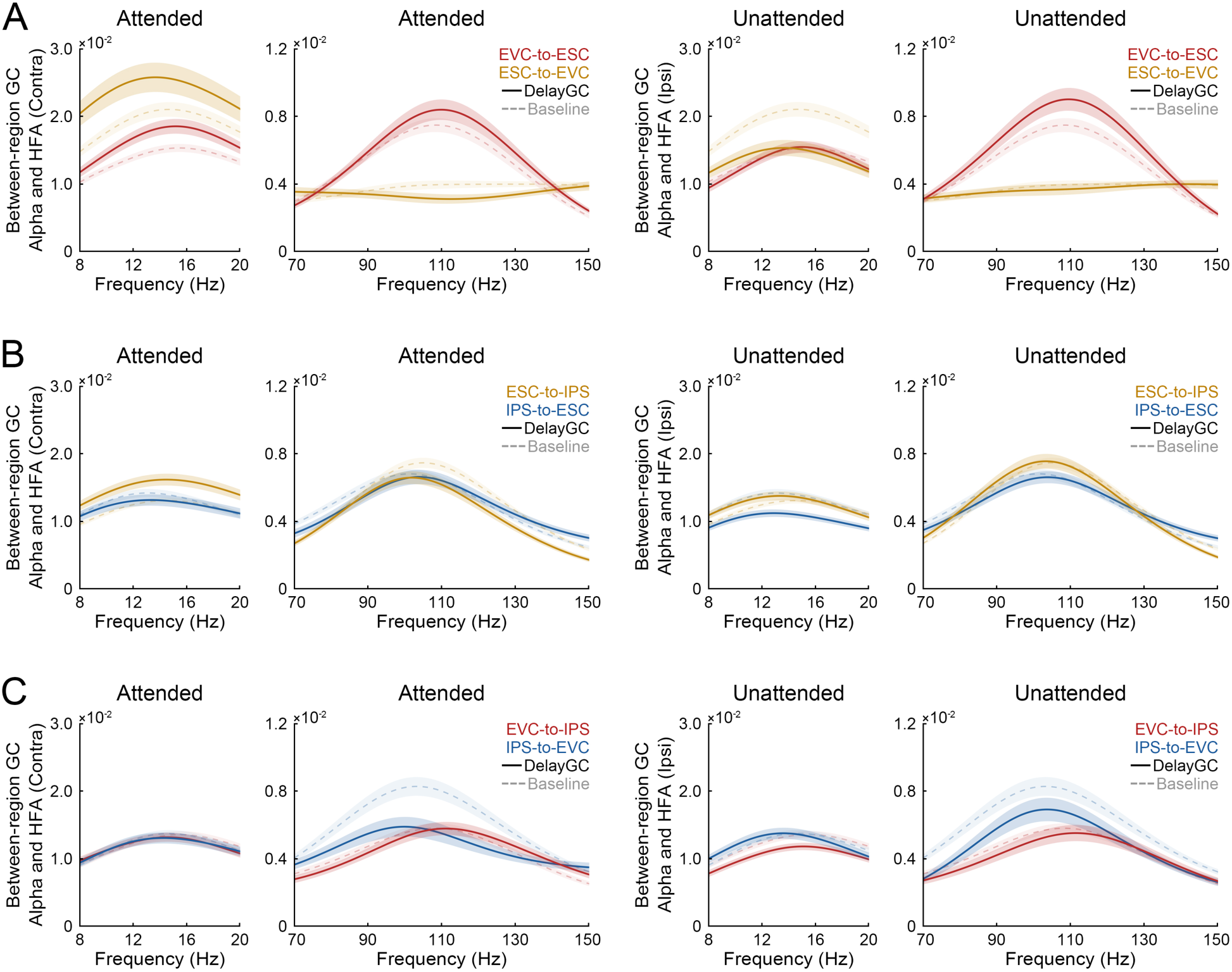
Pairwise Granger causality in visual topographic cortex. Granger causal (GC) influences of information flow along the visual hierarchy are illustrated for pairs of topographic areas at attended and unattended locations between EVC and ESC (**A**), between ESC and IPS (**B**), and between EVC and IPS (**C**), averaged across significant electrode pairs during the cue-target delay (solid line) and the pre-cue baseline (dashed line; shaded regions denote SEM). For each panel, the x-axis indicates alpha frequency band (8-20 Hz) and HFA frequency band (70-150 Hz), and the y-axis indicates Granger causal estimates. Note that the baseline GC influences measured at attended and unattended locations were combined to better estimate the baseline-level GC reference for each pair of visual topographic areas. All of these electrode pairs came from the four subjects (S1-S4) who had multiregional coverage of visual topographic areas.

**Table S1.** Electrodes with attention-modulated alpha responses in visual topographic and non-topographic cortex. One hundred and twenty-four electrodes were analyzed in visual topographic cortex from five subjects, including early visual cortex (V1-V3 dorsal/ventral: N = 37), extrastriate cortex (V3A/B, TO1-2, LO1-2 & VO1-2: N = 42), and parietal cortex particularly in areas along the intraparietal sulcus (IPS0-5 & SPL: N = 45). From all eight subjects, four hundred and forty-five electrodes in non-topographic cortex were analyzed for comparison purposes, distributing across parietal (N = 196), temporal (N = 131), and frontal lobes (N = 118). In order to quantify attention-modulated alpha responses in each cortical area, the numbers of task-responsive electrodes and contributing subjects (enclosed in parentheses) are summarized here, including electrodes that showed significant delay-related changes in alpha-band activity (‘Responsive’), electrodes that showed spatially selective changes in alpha-band activity (‘Selective’), and electrodes that showed significant within-region cross-frequency coupling (‘Coupled’) for the attended condition associated with alpha desynchronization (‘α_-_’), the unattended condition associated with alpha synchronization (‘α_+_’), and both conditions (‘α_-/+_’). Note that ‘Selective’ electrodes are included in ‘Responsive’ electrodes, and ‘α_-/+_’ electrodes are included in ‘α_-_’ and ‘α_+_’ electrodes.

| Cortical Regions | Total | Responsive |  |  | Selective |  |  | Coupled |  |  |
| --- | --- | --- | --- | --- | --- | --- | --- | --- | --- | --- |
| | | $\alpha_-$ | $\alpha_+$ | $\alpha_-/+$ | $\alpha_-$ | $\alpha_+$ | $\alpha_-/+$ | $\alpha_-$ | $\alpha_+$ | $\alpha_-/+$ |
| V1-V3 dorsal/ventral | 37(4) | 5(3) | 23(4) | 4(3) | 5(3) | 6(3) | 4(3) | 11(3) | 16(4) | 4(2) |
| V3A/B TO LO VO | 42(4) | 8(4) | 15(4) | 5(4) | 5(4) | 12(4) | 4(3) | 8(4) | 12(4) | 3(2) |
| IPS0-5 SPL | 45(5) | 22(3) | 9(4) | 3(1) | 9(2) | 3(1) | 3(1) | 14(5) | 16(4) | 4(2) |
| Parietal (non-topo) | 196(8) | 20(5) | 96(8) | 3(2) | 4(3) | 23(5) | 2(1) | 39(8) | 57(8) | 13(4) |
| Temporal (non-topo) | 131(6) | 5(4) | 88(6) | 2(2) | 2(2) | 9(4) | 2(2) | 38(6) | 47(5) | 20(4) |
| Frontal (non-topo) | 118(7) | 18(3) | 49(6) | - | 2(1) | 2(2) | - | 19(6) | 24(7) | 5(4) |
| Total Number | 569(8) | 78(7) | 280(8) | 17(6) | 27(6) | 55(7) | 15(5) | 129(8) | 172(8) | 49(8) |

